# A missense variant effect map for the human tumour suppressor protein CHK2

**DOI:** 10.1101/2024.02.13.579700

**Authors:** Marinella Gebbia, Daniel Zimmerman, Rosanna Jiang, Maria Nguyen, Jochen Weile, Roujia Li, Michelle Gavac, Nishka Kishore, Song Sun, Rick A Boonen, Jennifer N. Dines, Alexander Wahl, Jason Reuter, Britt Johnson, Douglas M Fowler, Haico van Attikum, Frederick P Roth

## Abstract

The tumour suppressor *CHEK2* encodes the serine/threonine protein kinase CHK2 which, upon DNA damage, is important for pausing the cell cycle, initiating DNA repair and inducing apoptosis. CHK2 phosphorylation of the tumour suppressor BRCA1 is also important for mitotic spindle assembly and chromosomal stability. Consistent with its cell cycle checkpoint role, both germline and somatic variants in *CHEK2* have been linked to breast and multiple other cancer types. Over 90% of clinical germline *CHEK2* missense variants are classified as variants of uncertain significance, complicating diagnosis of CHK2-dependent cancer. We therefore sought to test the functional impact of all possible missense variants in CHK2. Using a scalable multiplexed assay based on the ability of human CHK2 to complement DNA sensitivity of a *S. cerevisiae* lacking its ortholog *RAD53*, we generated a systematic ‘missense variant effect map’ for *CHEK2* missense variation. Map scores reflect known biochemical features of CHK2 and exhibit good performance in separating pathogenic from benign clinical missense variants. Thus, the missense variant effect map for CHK2 offers value in understanding both known and yet-to-be-observed CHK2 variants.

## Introduction

DNA lesions activate cell-cycle checkpoints, which are important for maintaining genome integrity[1,2]; [3]; [4]. *CHEK2*, a tumour suppressor gene encoding the serine/threonine checkpoint kinase 2 (CHK2), is an important checkpoint regulator of DNA repair, cell cycle regulation and apoptosis in response to DNA damage [5]. Germline variants in *CHEK2* have been linked to multi-organ cancer predisposition [6], and *CHEK2* is now typically included with *BRCA1*, *BRCA2,* and *PALB2* as a breast cancer risk gene [7,8]; [9].

Extensive sequencing has led to the identification of many *CHEK2* variants, both rare and common. Although *CHEK2* founder mutations c.1100delC and p.I157T (c.470T>C) have been shown to increase breast cancer risk by 2.6 and 1.4-fold respectively [8,10,11], the extent to which the vast majority of *CHEK2* variants are associated with elevated risk remains unclear. Indeed, 94.5% of the 1,519 *CHEK2* missense variants reported on ClinVar [12] are classified as variants of uncertain significance (VUS), limiting the value of genetic diagnosis of CHK2-dependent cancers.

According to current guidelines from the American College of Medical Genetics and Association of Molecular Pathology (ACMG/AMP), cell-based or *in vitro* functional assays of variant impact are one of the stronger forms of evidence for clinical variant interpretation. Although functional assay results are typically unavailable for rare variants, mutational scanning technologies have made it possible to systematically test variants at a large scale [13,14]. The resulting variant effect (VE) maps, measuring the functionality of nearly every possible coding variant, can provide evidence ‘proactively’ (even in advance of the first clinical presentation of a variant). A systematic evaluation of clinical missense variants in tumour suppressors *TP53*, *BRCA1, PTEN,* and *MSH2* recently found that, of variants that were previously annotated as VUS and had VE map evidence, more than half could be more informatively classified [15]; [16]

To assess the functional consequences of all possible mutations in the *CHEK2* gene, we exploited a yeast-based complementation assay. *CHEK2* is the human homolog of *Saccharomyces cerevisiae RAD53*, which also helps to maintain genome integrity after DNA damage [17]. Yeast strains lacking *RAD53* lose viability and are unable to recover from genotoxic stress [18]. Despite one billion years of divergence between yeast and humans, yeast functional complementation assays have been shown to detect ∼60% [19] of pathogenic variants at a stringency where 90% of variants identified as damaging are pathogenic [19–22]. Wild-type human CHK2 restores the loss of Rad53 activity after DNA damage is induced in the presence of methyl methanesulfonate (MMS). This complementation relationship, previously exploited at smaller scale [23], offers a facile assay to test the functional impact of CHK2 missense variants.

We integrated this functional assay within a deep mutational scanning framework [24][13] to produce a comprehensive functional VE map of human CHK2 in the presence of MMS. We validated this map based on agreement with known biochemical features of CHK2, smaller-scale functional assay results and the ability to separate pathogenic and benign variants.

## Results

To assess the impact of human CHK2 variants in response to DNA damage, we carried out functional assays for nearly all possible missense variants (Fig. 1A). Below, after confirming validity of a previously published assay, we describe the large-scale variant testing process in greater detail, then relate the resulting functional scores to known biochemical and structural features of *CHEK2*, and to clinical variant pathogenicity.

**Fig. 1.**
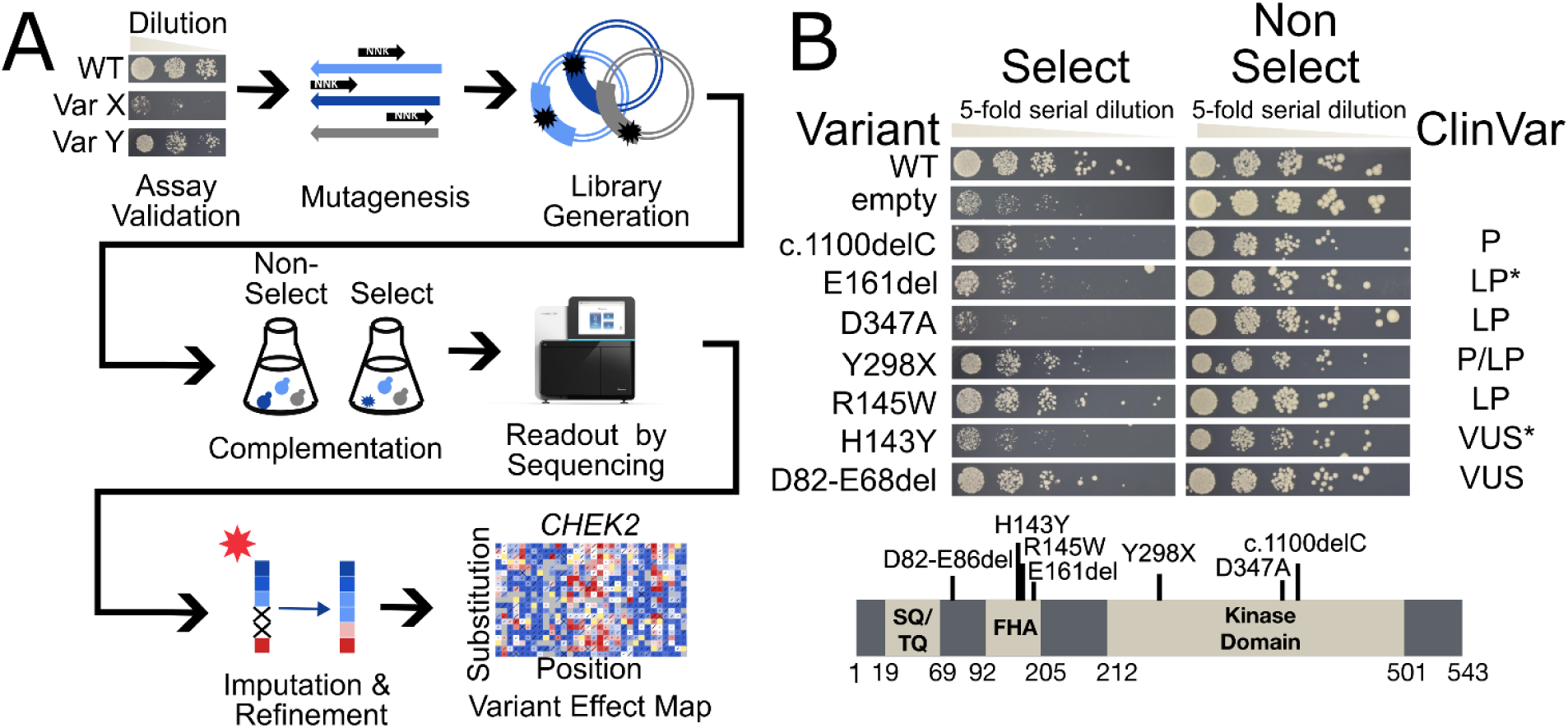
Assay validation and generation of the *CHEK2* variant effect map. **A** Overview of the TileSeq framework that was followed to produce the *CHEK2* variant effect map. The red star indicates the optional imputation and refinement stage of the process. **B** Yeast-based functional complementation assay for CHK2 domains labelled with tested variants. The *sml1Δrad53Δ* yeast strain was transformed with the expression vector pADH1-Leu carrying wild-type (WT) *CHEK2* or the empty pADH1-Leu expression vector (empty). The selective condition was 0.007% MMS and the non-selective condition was 2% DMSO as a positive control. Yeast growth was assessed by spotting serial dilutions of yeast cells on selective media and incubation for three days. P, LP, and VUS indicate pathogenic, likely pathogenic, and variant of uncertain significance, respectively. We note LP* for variant p.E161del because it has been annotated variously as LP, P, and VUS, and note VUS* for variant p.H143Y because 1 of 6 annotations in the ClinVar database was LP, while the others were VUS.

### Confirming and validating a yeast complementation assay for human CHEK2

Yeast is dependent on *RAD53* for both viability and response to stress [18], and for resistance to DNA damage induced by MMS [25,26]. Building on observations that expression of human CHK2 in yeast rescues the loss of *RAD53* [18], and on the observation that *sml1*Δ *rad53*Δ yeast strains are viable but sensitive to DNA damage [27,28], Roeb *et al.* measured the ability of dozens of CHK2 variants to rescue the MMS sensitivity phenotype of an *sml1*Δ *rad53*Δ yeast strain [23], and this assay was later used to test over 100 CHK2 missense variants [23]. Here we recapitulated the complementation assay of Roeb *et al.* with minor modifications (see Methods) and showed for a set of control variants that (with the single exception of pathogenic variant R145W) we observed the expected impacts on MMS sensitivity (Fig. 1B).

### A proactive missense variant effect map for CHK2

To exploit the yeast-based complementation assay at scale, we adopted the TileSeq framework [13] (see Methods), using the following steps:

First, we generated variant libraries of *CHEK2* missense variants by Precision Oligo-Pool-guided Code alteration (POPCode) mutagenesis [13], with each library focused on one of four CHK2 regions (Fig. S1), each ∼150 amino acids in length. Within each mutagenized region we sequenced short (∼151 nt) ‘tile’ segments which collectively span the region (see Methods).

Second, we used recombinational cloning to transfer each of these mutagenized *CHEK2* amplicon libraries *en masse* into the appropriate yeast expression vector, then individually transformed each vector pool into *sml1*Δ *rad53*Δ yeast and stored frozen aliquots of each transformed yeast pool (“pre-selection pools”).

Third, to obtain post-selection pools we grew yeast pools on solid media with 2% glucose and 0.007% MMS to select for cells expressing functioning CHK2 variants.

Fourth, variant frequencies were obtained by sequencing ∼2M or more reads for each *CHEK2* tile from each yeast pool from both pre- and post-selection yeast pools, and also from a control WT sequence library.

Clones in the pre-selection pool were estimated to contain an average of 0.3 amino acid changing variants per clone, with less than 5% of clones bearing multiple variants (Table S1). Estimates of variant frequency in the pre-selection library were strongly correlated between sequencing libraries from replicate cell pools (Pearson Correlation Coefficient or PCC = 0.98; P < 1e-320 (Fig. S2A). To score the impact of selection on each variant, we measured the log-ratio of variant frequency (log(ɸ)) in each post-selection pool relative to the corresponding pre-selection pool (see Methods). To remove less well measured scores, e.g., where pre-selection abundance was too low, we applied filtering criteria (see Methods). Replicate log(ɸ) scores were significantly correlated (PCC = 0.64; P <1e-320) (Fig. S2B). Log(ɸ) scores in each region were next calibrated to yield more interpretable functional scores such that a functional score of one corresponds to the mode of synonymous variants, and a functional score of zero corresponds to the mode of nonsense variants (excepting positions near the C-terminus as discussed below).

Nonsense variants up to position 489 exhibited highly damaging scores (mean score of 0.0 +/- 0.5 SD). However, nonsense variants from position 490 to the C-terminus appeared less damaging (mean score 0.7 +/- 0.5 SD), suggesting that these truncating protein variants are functional (although these nonsense variants may still be damaging as noted in the Discussion)(Fig. S3). After excluding nonsense variants beyond position 489, nonsense and synonymous variants showed distinct functional score distributions (Fig. S4).

Together, this process yielded a missense variant effect map with functional scores for 7955 amino acid substitutions and 419 nonsense mutations (Fig 2 and Fig. S5A), corresponding to over 77% of all possible amino acid changes (Fig. S6). Most missense variants appeared ‘synonymous-like’ (∼64.5% of variants scored above 0.5), while a substantial proportion of variants yielded ‘nonsense-like’ functional scores, with ∼25% of variants having scores below 0.2. For *CHEK2*, 3182 amino acid substitutions are achievable by a single-nucleotide variant (SNV) and thus more likely to be observed in humans, and our map covered 2837 (89%) of these (Fig. S6).

**Fig. 2.**
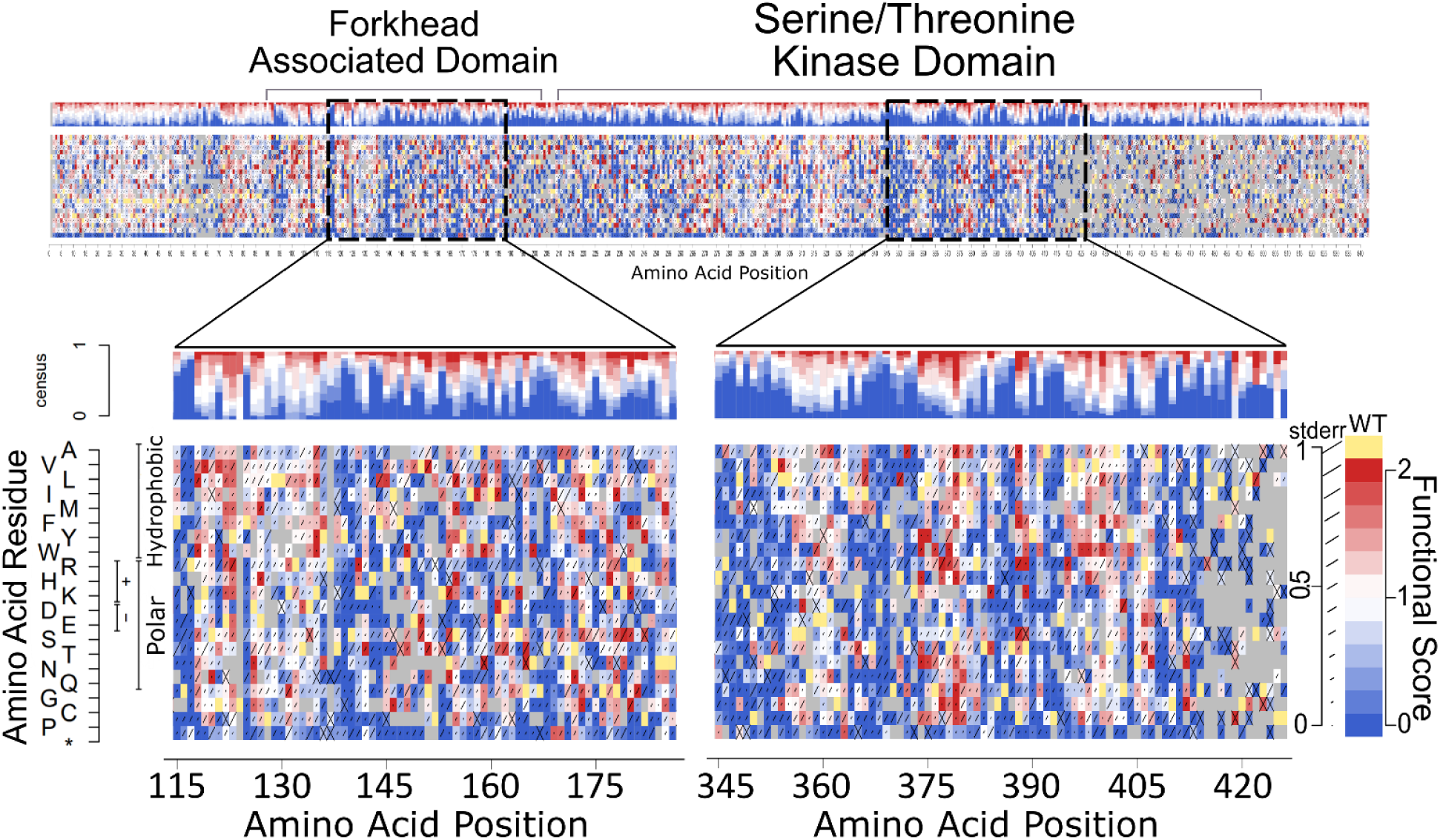
Experimental variant effect maps of CHK2. A preview of the full-length original CHK2 variant effect (VE) map with magnified views of segments of the FHA domain (positions 116-186) and kinase domain (positions 346-426). An enlarged version of the map can be found in Supplementary (Fig. S5A). Each heatmap shows functional scores for every possible amino acid substitution and nonsense mutation (bottom row). Above each heatmap is a consensus summary of the distribution of functional scores at each position. As shown in the legend (right): A functional score of 0 (blue) corresponds to the median score of nonsense variants; a score of 1 (white) corresponds to the median of synonymous variants; scores above 1 (red) are considered hyper-complementing; grey indicates missing data; and the length of the diagonal line on each block indicates estimated each score’s estimated standard error.

It has been previously shown that, given variant effect data for a sufficient number of substitutions at a given amino acid position, missing values can be imputed with accuracy approaching that of experimental measurements. Because imputation is based only on information intrinsic to the TileSeq experiment, this remains independent of other computational variant effect predictors (see Methods). Variant effect maps can then be refined by averaging the original and imputed values [13]. Of the 2343 substitutions missing from the map, we were able to impute 1441 (61.5%) substitutions (see Methods). For the 7955 amino acid substitutions with well-measured scores, 7206 (90.6%) were refined (see Methods), with a limited fraction of variants shifting substantially (e.g., 253 (3.5%) of refined variants either shifted from 0.8 to below 0.2 or from below 0.2 to above 0.8 after refinement). After generating an imputed/refined version of the map (Fig. S5B), we saw that the imputation and refinement process slightly improved separation of missense variant scores (Fig. S4A and Fig. S4C). However, because the impact of the imputation/refinement process on our results was modest, our analysis here is focused on the original map scores.

### Functional scores reflect known structural and biochemical features of CHK2

Scores in our variant effect map generally agreed with conservation and known structural/biochemical features of the CHK2 protein. For example, lower-scoring variants indicated in dark blue (Fig. 2) occurred more commonly at conserved positions (Wilcoxon rank-sum test, |Δ median| = 0.35, p = 1.97e-10) (Fig. 3A). As expected, solvent-accessible residues scored higher than non-accessible positions in the FHA (amino acids 92 - 205) and kinase domains (amino acids 212 - 501), as well as across the entire protein (Wilcoxon rank-sum test yielded |Δ median| values of 0.39, 0.28, 0.23, respectively, with p = 0.004, 1.3e-5, and 3.4e-7, respectively) (Fig. 3B). As one example, the founder variant p.Ile157Thr (allele frequency ∼1.1%), which is buried at the centre of the hydrophobic core, showed a low functional score for most polar substitutions but not for hydrophobic substitutions. Visualizing the dimeric CHK2 crystal structure (PDB 3i6u) with each residue coloured according to the median map score for each position further supports the expected finding that the map tended to give more damaging scores to buried residues (Fig. 3C and Fig. 3D).

**Fig. 3.**
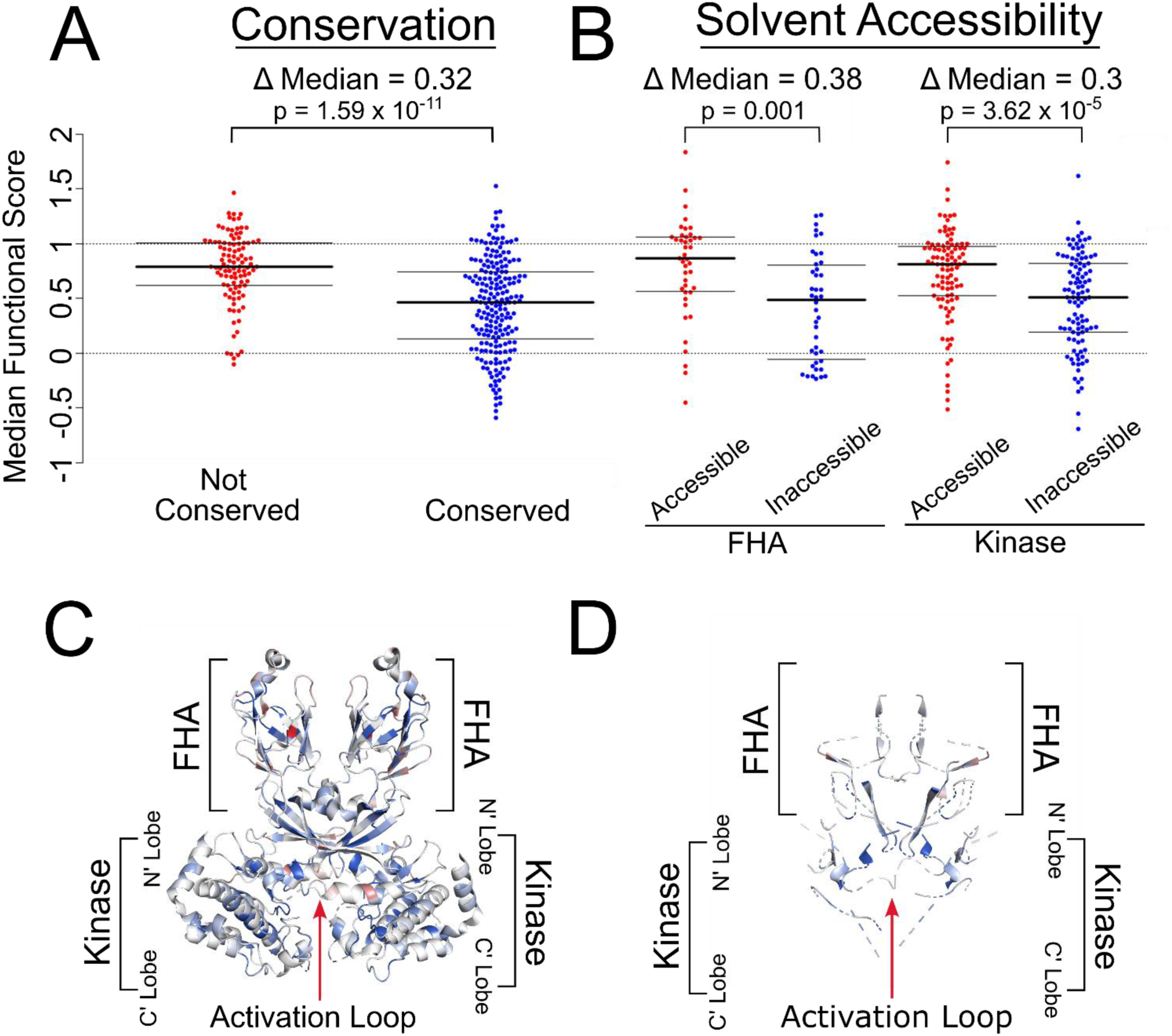
*CHEK2* map scores agree with expected effects of conservation and solvent accessibility. **A** Median *CHEK2* functional scores were calculated and stratified based on each position’s conservation score. Conservation scores were derived from ConSurf multiple sequence alignments based on an Alphafold predicted PDB file for CHK2. Residues labelled ‘Not Conserved’ have ConSurf scores above 0.91 while ‘Conserved’ positions are below −0.47. **B** Solvent Accessibility was determined using FreeSASA software [version 2.02 October 22^nd^, 2017] on the AlphaFold [version 2022-11-01, Monomer v2.0 pipeline; UniProt O96017] *CHEK2* structure. Residues with less than 20% accessible surface area (ASA) for their side-chain were considered ‘Inaccessible’ while those with above 50% side-chain ASA are ‘Accessible’. Results were separated into the FHA domain (positions 92-205) and Kinase domain (positions 212-501). **C** Crystal structure of the CHK2 dimer with coloured residues based on the median functional score of each position. **D** Crystal structure of the CHK2 dimer with blue residues for positions with a dimer interface burial greater than 0%.

Stop codon variants appeared damaging up to amino acid position 489 but were generally tolerated afterwards, suggesting that proteins truncated after this point can carry out their function (Fig. S3). We note, however, that our assay is not expected to capture impacts of nonsense-mediated decay that might occur in the endogenous context. Positions within the kinase domain exhibited significantly lower functional scores than the rest of the protein (Wilcoxon rank-sum test showed |Δ median| = 0.11, with p = 1.6e-4). Although positions within the shorter FHA domain showed a numerically similar reduction in functional score, this result was not significant (Wilcoxon rank-sum test |Δ median| = 0.11, with p = 0.21) (Fig. S7).

We next compared *CHEK2* functional scores with computational predictions of variant impact on protein stability. Our functional scores showed significant correlation with predicted change in folding energy (ΔΔG from DDGun [29]; PCC = 0.4; P = 1.2e-22), further validating our variant effect map. It has been previously noted that positions where substitutions impact overall functionality, but not stability, may correspond to positions that are important for catalysis, regulatory modification, structural flexibility, or physical interaction with other biomolecules [30]; [31]; [32]. Evaluating agreement between functional and ΔΔG scores within successive sequence ‘windows’ revealed five regions in the FHA domain where substitutions impacted function more profoundly than stability (Fig. 4), including residues 93-97, 115-117, 138-144, 164-169, and 191-202. Because the FHA-FHA dimerization interface is reportedly centred on the aromatic side chains of Trp97 and Phe202, and because dimerization is required for CHK2 activation, it is likely that all of these positions are important for dimerization but not for stability of the monomer. Additionally, Ser140, which is required for dimer dissociation after activation, has minimal impact on protein stability despite low functional scores [33,34]. Excluding these five regions further increased the correlation between functional and ΔΔG protein stability scores (PCC = 0.45) (Fig. S8).

**Fig. 4.**
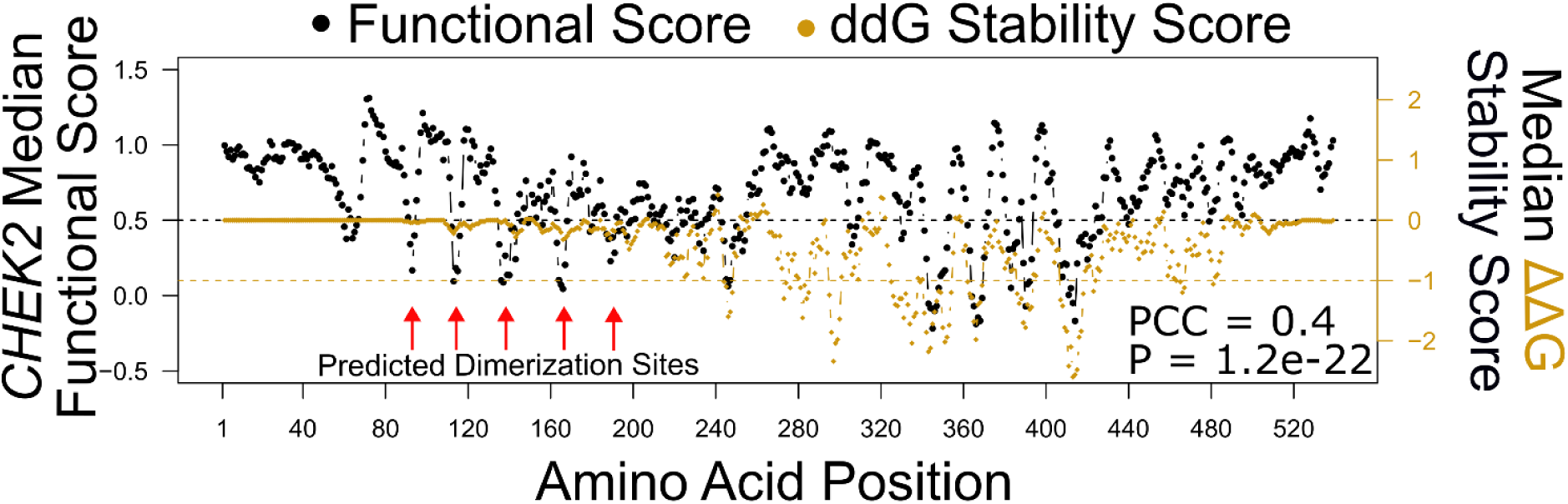
Regions with functional but not (predicted) protein stability impacts highlight positions in the FHA domain important for dimerization. Stability scores (estimated ΔΔG folding energy values) were generated via the DDGun software using only the CHK2 protein sequence as input. At each position, ΔΔG was calculated for every substitution and the mean value was plotted against the average positional CHK2 functional scores in a five amino acid wide moving window analysis (with single-residue step size). Variants at positions with a ΔΔG near 0 are predicted to be typically stable while scores less than −1 are predicted to be typically unstable. CHK2 functional scores of 1 are wildtype-like while scores near 0 are considered to have profound loss of function. The dashed horizontal black line indicates a detrimental functional score of 0.5 and the dashed horizontal gold line an unstable ΔΔG score of −1.

CHK2 activation requires phosphorylation of Thr68 in the SQ/TQ cluster domain (SCD) by DNA damage-activated ATM kinase. It has been shown that ATM-induced phosphorylation causes transient dimerization of CHK2, thereby inducing kinase activation through autophosphorylation of Thr383 and Thr387 [33,34]. Consistent with this model, phosphomimetic mutation T68E shows near-wildtype function with a map score of 0.8 while the phospho-dead variant T68A shows complete loss of function with a score of −0.6 (Fig. S9). Positions critical for dimerization, Thr68 and Ile157 [10,35], as well as the catalytic residue Asp347 and phospho-acceptors Thr383 and Thr387 all have low median scores of 0.2, 0.4, −0.6, 0.0, and −0.6 respectively. The activation loop, corresponding to positions 371-391 in the kinase domain, displayed variable tolerance to substitutions: eleven residues appeared highly tolerant to variation (with median scores greater than 0.8) while seven positions, including Thr383 and Thr387, had median functional scores less than 0.2. Because auto-phosphorylation accompanying CHK2 activation requires ATP-binding via hydrogen bonding and hydrophobic interactions [4,36], we examined residues that had been previously identified as important for CHK2 activation on the basis of CHK2 crystal structures, based on proximity to either complexed ADP or the competitive ADP inhibitor debromohymenialdisine (DBQ) [34] (Table S2). Of these 16 residues, 13 (81%) exhibited map scores with median scores below 0.5.

### CHK2 map scores correlate with mutational hotspots in the kinase domain

In a previous study, the combination of multiple sequence alignments of candidate tumour suppressors and missense mutation rates at each position were used to identify mutational hotspots [37]. Some of the 23 mutational hot spots identified overlap with known protein features. For example, positions 247 and 248 in the VAIK motif, as well as position 368 in DFG, are located in ADP binding sites and the APE-6 hotspot is located in the kinase activation T-loop at position 386. Of the 23 CHK2 hotspots identified, all 16 that fell within a named motif (APE, VAIK, HRD, DFG and the three G positions within the GxGxxG motif) scored as intolerant to variation in our map (median scores below 0.5). Median values of the hotspot positions within the VAIK, HRD, and APE motifs located at positions 246-249, 345-347, 392-394, respectively, exhibited damaging scores ranging from −0.76 to 0.35 (Table S3). Within the GxGxxG motif, hotspot positions G227, G229, and G232 had damaging median scores of 0.19, 0.06, and −0.35 respectively, while hotspot positions corresponding to the DFG motif medians were −0.62 (D368), −0.42 (F369) and 0.07 (G370). These 16 mutational hotspots are evidently important for CHK2 function as demonstrated by scores that were significantly lower than CHK2 overall (Wilcoxon rank-sum test, |Δ median| = 0.95, p = 2.0e-41) (Fig. 5). Of the six additional hotspot positions identified by Hudson *et al.* [37] but not listed above, only three appeared damaging in our map: HRD+5 (N352), HRD+7 (L354), and APE-6 (G386) showed median scores of 0.04, −0.22, and 0.05 respectively.

**Fig. 5.**
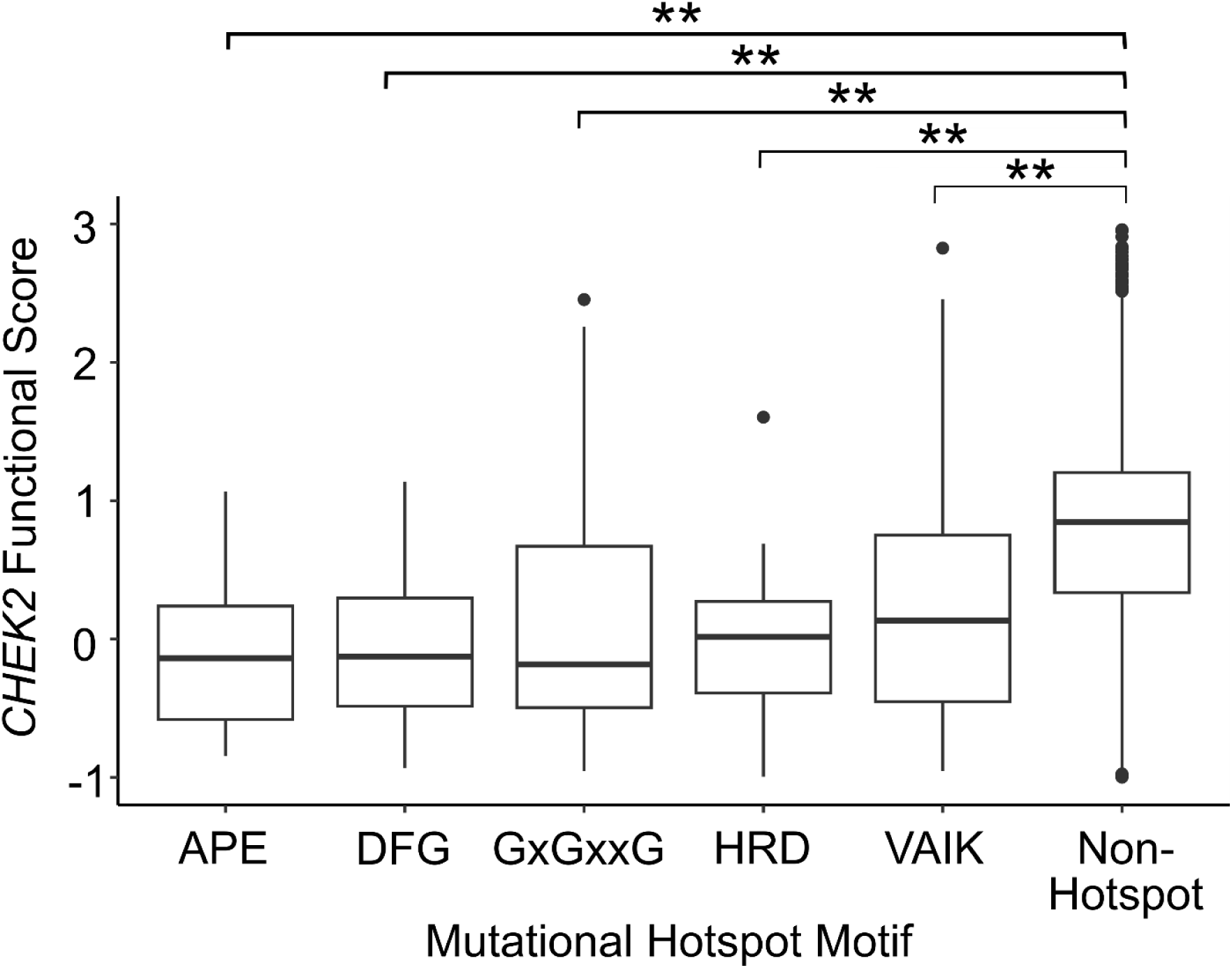
Known mutational hotspots in CHEK2 are largely intolerant to missense variants. Mutational hotspots identified in *Hudson et al.* [37] were separated into distinct kinase activation motifs (APE, DFG, GxGxxG, HRD, and VAIK) and plotted against *CHEK2* functional scores for each variant at these positions. Boxplots depict the median functional score for each motif as well as the 25th and 75th percentiles with vertical lines extending to 1.5 times the interquartile range above and below 25th and 75th percentiles. All motifs were conserved, matching the consensus sequence observed in other kinases. Double stars indicate significance by Mann Whitney U test of less than 1e-5 as compared to non-hotspot positions. The absolute delta medians for motifs APE, DFG, GxGxxG, HRD, and VAIK, as compared to non-hotspot positions, are 1.07, 1.08, 1.04, 0.96, and 0.81 respectively.

### Relating the CHK2 map to previously-reported functional assays

The functional effects of CHK2 missense variants have been previously assessed in a variety of assays that include rescue of growth (of yeast cells) under DNA damage [11]; [23], and mammalian cell-based protein stability and kinase activity assays [9]; [38]. Our map scores showed modest but significant correlation with the Delimitsou *et al*. yeast growth-based study (ref. [11], R = 0.44, p = 2.1e-6), but no significant correlation with the Roeb *et al*. (ref. [23], 2012; R = 0.19, p = 0.38). Variants classified as “functional” in Delimitsou *et al*. had a median score of 0.95 in our map, and variants deemed “non-functional” had a median score of 0.34 (Fig. 6A).

**Fig. 6.**
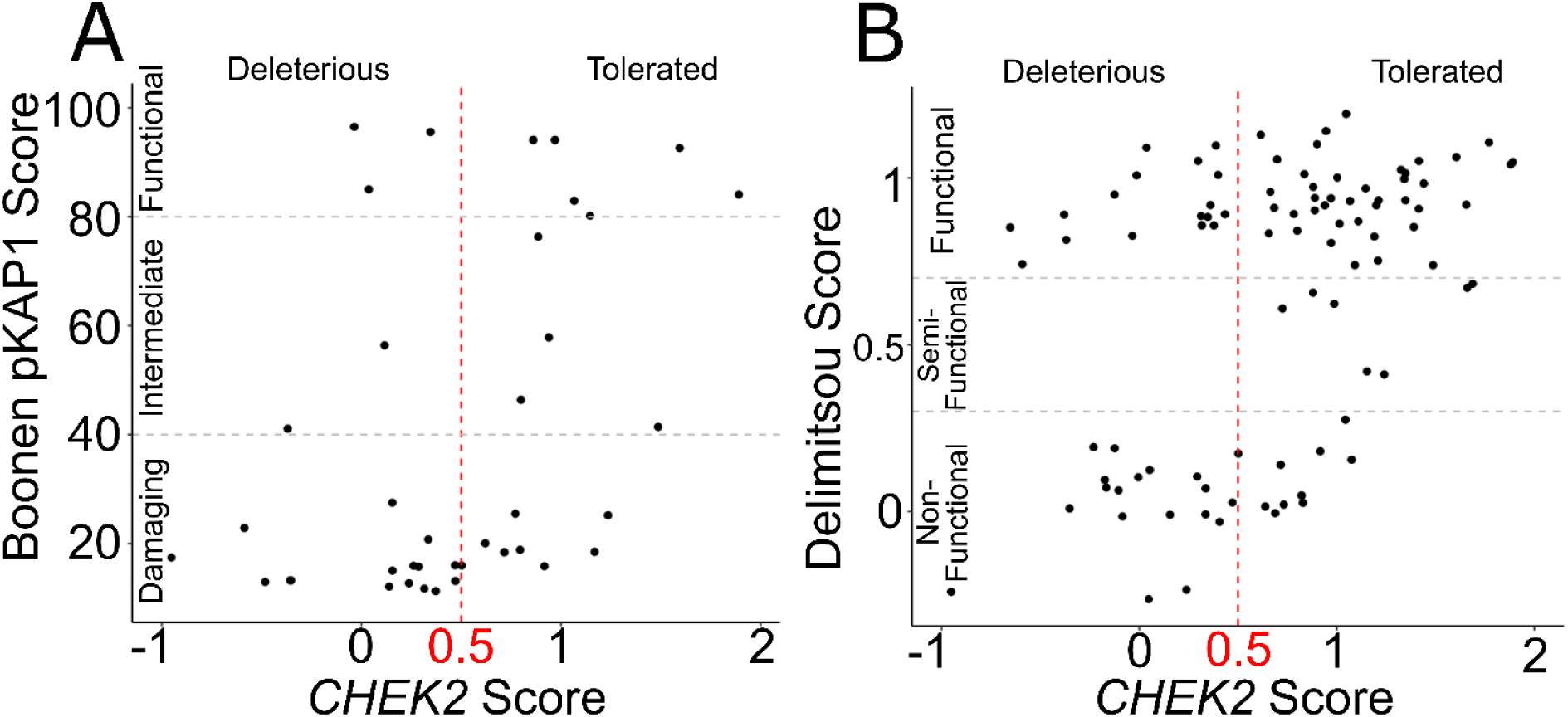
*CHEK2* variant effect map scores agree with mammalian and yeast functional assays. Scatterplot relating *CHEK2* variant effect map scores (*x*-axis; indicating map score threshold of 0.5 for reference) to: **A** measurements of pKAP1 phosphorylation after ionizing radiation in mouse embryonic stem cells (*y*-axis), indicating previously assigned categories of damaging, intermediate, and functional scores [9]; and **B** functional complementation scores for 120 *CHEK2* missense variants in yeast in the presence of MMS (*y*-axis; indicating map score threshold of 0.5 for reference), indicating previously assigned categories of non-functional, semi-functional, and functional [11].

We also showed modest but significant correlation with assays by Boonen *et al.* of the ability of CHK2 to phosphorylate the downstream substrate KAP1 in mammalian cells (ref. [9], R = 0.41, p = 0.007) (Fig. 6B). Variants classified as “functional” by Boonen *et al.* had a median score of 0.97 in our map, in contrast to a median map score of 0.32 for the variants they called “damaging”. We observed a low (and not significantly different from zero) correlation coefficient between our map scores and another mammalian cell based study [38] (R = 0.30, p = 0.16) of pKAP phosphorylation and the Boonen *et al*., 2022 protein stability assay (R = −0.09, p = 0.66) (Fig. S10).

In a more detailed comparison of our map with the two published assay sets showing significant correlation with our map– the Boonen *et al.* pKAP1 assay and the Delimitsou *et al.* yeast growth assay, we examined the 18 clinical missense variants tested in all three studies finding that 9 had results that agreed in all three. All assays correctly detected many variants as damaging, e.g p.D347N, a position known to impact kinase function [39], and residues associated with elevated breast cancer risk such as p.Y390S [12][40]. Interestingly, both p.D347N and p.Y390S had conflicting clinical annotations, with a mixture of either “likely pathogenic” or “pathogenic” and “VUS’’ annotations. In positions adjacent to Asp347 and Tyr390, all three assays found detrimental effects for p.R346H and p.A392V (with predominantly VUS annotations). Predicted dimerization variants p.R117G and p.G167R, which exhibited damaging scores in all three studies, are predominantly annotated as likely pathogenic or pathogenic. Only three variants – p.R180C, p.R181H, and p.N446D – were found to have normal function in all assays. ClinVar describes a mixture of likely benign or benign and VUS clinical annotations for all three variants, with one submitter annotating p.R180C as likely pathogenic. Thus, our map, taken together with previous assay data, concurs with pathogenic clinical annotations for p.R117G and p.G167R, adds to the weight of evidence towards pathogenicity for p.R346H, p.D347N, p.Y390S and p.A392V; and provides evidence towards benignity for p.R180C, p.R181H, and p.N446D.

Of the remaining nine variants for which results differed between our map and either the Boonen *et al*. pKAP1 assay or the Delimitsou *et al*. yeast growth assay, three substitutions (p.D203G, p.E239K, and p.D438Y) had intermediate scores in Boonen but were seen as well-tolerated by Delimitsou *et al.* and our map. Conversely, some variants with intermediate scores in our assay were observed to be damaging (p.I160T, p.D162G) by both Boonen *et al*. and Delimitsou *et al*. studies or found to be non-damaging (p.I157T) by both. Finally, our assay found p.R145W to be well-tolerated and variants p.C243R and p.N186H to be damaging, while Boonen *et al*. and Delimitsou *et al.* both found results opposite to ours for each of these. All nine of these variants except R145W (which was classified as likely pathogenic in ClinVar) either had conflicting interpretations or were classified as VUS.

#### *Ability of the* CHK2 *map to distinguish pathogenic from benign variation*

We evaluated the ability of our CHK2 map to distinguish pathogenic from benign variants. Reference annotations for 21 pathogenic and 39 benign missense variants were independently derived by Invitae using Invitae’s previously published Sherloc variant classification framework [41] [42]. Of these, our *CHEK2* variant effect map provided scores for 20 pathogenic and 36 benign variants.

We examined performance in terms of precision (fraction of variants with scores below a given threshold that are known to be pathogenic) and recall (fraction of known pathogenic variants that received a score below this threshold). Because precision is dependent on the (somewhat arbitrary) balance of pathogenic and benign variants in the reference sets, we instead calculated balanced precision values (precision expected with a reference set containing 50% pathogenic and 50% benign variants). Our map achieved an area under the balanced precision vs. recall curve (AUBPRC) of 0.8, for example detecting >70% of pathogenic variants at a balanced precision of >80% (Fig. S11).

We next wondered whether performance in distinguishing pathogenic from benign variation could be enhanced by refraining from making predictions for variants with less certain scores. To this end, we eliminated variants for which the confidence interval encompassed a midpoint threshold of 0.5. For the 12 pathogenic and 28 benign variants remaining after this step, AUBPRC improved to 0.87, with 33% of pathogenic variants detected at a stringency achieving 100% balanced precision and 50% recall at 90% balanced precision, while still detecting >70% of pathogenic variants at a balanced precision of >80% (Fig. 7A).

**Fig. 7.**
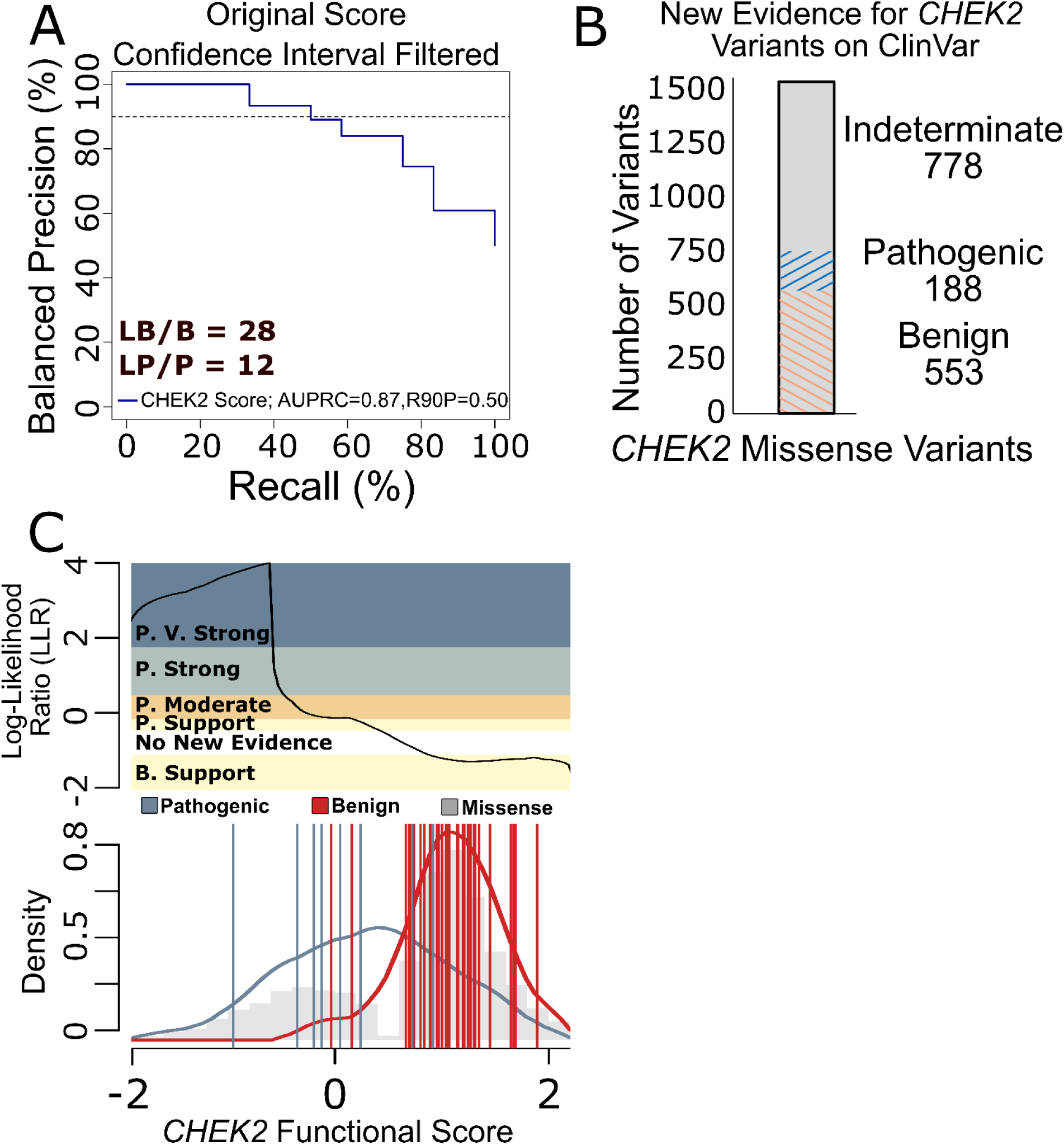
*CHEK2* functional scores correctly classify variants in ClinVar with known annotations and provide evidence for VUS reclassification. **A** Using *CHEK2* map scores after confidence interval filtering and a known set of clinically annotated pathogenic or benign *CHEK2* variants from Invitae, we evaluated balanced precision—defined at each score threshold by the fraction of variants that are pathogenic given a balanced (50% prior probability of pathogenicity) test set—versus recall (fraction of pathogenic variants captured at this threshold). The horizontal dashed line indicates R90BP with the numerical AUPRC and R90BP listed in the bottom-left hand legend. LB/B indicates likely benign and benign while LP/P means likely pathogenic and pathogenic. **B** *CHEK2* functional score ranges were converted into ACMG evidence strengths using Log-Likelihood Ratios (LLR) and matched against all VUS in ClinVar (as of June 2023). Evidence towards pathogenicity or benignity are split into discrete categories where V. Str, Str., Mod., and Sup., mean very strong, strong, moderate, and supporting respectively. Sections in blue represent evidence towards pathogenicity, red towards benignity, and grey are VUS where no new evidence is provided. The numbers below to each label show how many variants fall into each category. **C** LLRs of pathogenicity were calculated by comparing the probability distributions of scores for known pathogenic and benign variants in the reference set. The log ratio between likelihood of observing a score in the positive pathogenic reference set (blue) compared to the negative benign reference set (red) was calibrated to ACMG evidence strengths [43]. Probability distributions are overlaid on the grey histogram of *CHEK2* missense variant scores with the top panel showing which score ranges correspond to each ACMG evidence strength.

We next wished to calibrate the value of *CHEK2* map scores under current clinical variant classification. Using the set of reference variants derived from filtering the original *CHEK2* scores, we first calculated Log-Likelihood Ratios (LLR) of pathogenicity for every variant’s functional score, by comparing the score distributions observed for pathogenic and benign variants (see Methods). Conversion from LLR values into evidence strengths as described in ACMG/AMP guidelines was done using an adaptation [32] of the Tavtigian *et al.* approach [43] (see Methods). This calibration assigned “very strong” evidence of pathogenicity for LLR scores above 2.5, “strong” evidence for scores between 2.5 and 1.3, “moderate” evidence for scores from 1.3 to 0.64, “supporting” evidence of pathogenicity for scores from 0.64 to 0.32. It further assigned “supporting” evidence of benignity for LLR scores between −0.32 and −1.32. This calibration of evidence strength aligns with the medians of nonsense and synonymous variants where nonsense-like scores around 0 suggest pathogenicity while synonymous-like scores of 1 suggest benignity. Based on the calibration, the CHK2 map provided at least a supporting level of evidence for 741 out of 1519 *CHEK2* VUS in ClinVar (Fig. 7B), including 30, 105, 9, and 44 VUS for which the map provided supporting, moderate, strong, and very strong evidence for pathogenicity, respectively, and 553 VUS for which the map provided supporting evidence of benignity (Fig. 7C).

## Discussion

The missense variant effect map we generated for CHK2, covering >77% of all possible amino acid changes and 89% of SNV-reachable amino acid changes, both recapitulates many known biochemical features of CHK2 and offers a new window on sequence-structure-function relationships and potential clinical applications.

### Correspondence to known biochemical features of CHK2

As expected, our map tended to show low (damaging) functional scores for conserved positions (Fig. 3A). Also as expected, solvent-accessible residues appeared more tolerant to variation than non-accessible positions across the entire protein, especially in the FHA (amino acids 92 - 205) and kinase domains (amino acids 212 - 501) (Fig. 3B). Damaging variants were also enriched in known mutational hotspots, with median scores near 0 for substitutions in the reported hotspot motifs–APE, DFG, GxGxxG, HRD, and VAIK (Fig. 5).

### Identifying residues important for CHK2 homodimerization

Variant effect maps can provide new hypotheses about the importance of specific amino acid residues. One approach to this is to combine scores and proximity of residue positions to known protein features. For example, we note that Tyr220 in the N lobe of the kinase domain appeared intolerant to variation (median score 0.08). Although its role is not well described, Tyr220’s proximity to Ile221 and Lys224–which are contact sites of the homodimerization interface between CHK2’s FHA and kinase domains [33] coupled with Tyr220’s low map scores, suggests that Tyr220 serves to stabilise the FHA/kinase dimer. The FHA-kinase dimerization (FHA-KD) interface is mediated by contacts between hydrophobic residues from both the FHA domain (Gly151, Pro152, Ile157, Tyr159, and Pro182) and the N lobe of the kinase domain (Ile221, Leu226, Leu236, Phe238, Cys243, Lys224, and Lys245) [33]. The results from our map support the idea that close proximity of Tyr220 to the hydrophobic core of the FHA-KD interface is important for maintaining favourable contacts and stabilising the FHA-KD interface.

A similar logic also suggests a role for kinase domain position Ala237 in stabilising the FHA-KD interface. With a low median score of 0.2, Ala237 is also located at the FHA-KD dimerization interface and is in close proximity to Phe238 which, along with Leu236 and Lys224, is reported to make van der Waals contact with Ile157 in the FHA-kinase domain dimerization interface [33]. The substitution of non-polar Ala237 with polar or bulkier amino acids could perturb contacts in the hydrophobic core and thus affect FHA-KD dimer stability.

Several cancer-associated germline variants at the FHA-KD interface [45] are classified as variants of uncertain significance [12] but have low map scores, suggesting their importance. These include: A247D (map score 0.2), a highly conserved residue in the kinase domain in close proximity to Cys243 and Lys245 in the hydrophobic centre of the FHA-KD interface which has previously been reported to affect protein function [46]; P152S (map score 0.2), located at a β5’- β6’ hairpin very close to Ile157 in the hydrophobic core of the interface, which could alter the hairpin conformation and destabilise the dimer by affecting the van der Waals contacts [33]; and R181C (map score 0.3), located at the periphery of the FHA-KD interface, which could affect intramolecular contacts and cause instability at the interface [33].

During CHK2 activation, a second homodimerization interface is formed between the FHA domains of the two protomers [47], mediated by van der Waals and hydrogen bond contacts from three adjacent β strands. The FHA-FHA interaction is mainly centred around the aromatic side chains of Trp97 and Phe202 which make contacts with Trp97 and Phe202, respectively, in the homomeric partner [33]. Substitution at these positions or in positions near the FHA-FHA interface may interrupt van der Waals and hydrogen bond contacts and further affect the dimerization [33]. Supporting this idea, positions adjacent to Trp97 (93-96) have a median score of 0.0 in our map indicating abnormal function.

As described in Results, the comparison of measured impacts on function with predicted impacts on stability (ΔΔG) can reveal active site residues or other roles beyond providing stability. For CHK2, this analysis revealed five regions in the FHA domain (positions 93-97, 115-117, 138-144, 164-169, and 191-202) where substitutions impacted function more profoundly than stability (Fig. 4). It is interesting to note that in the region 138-144, in which substitutions appear to impact protein function but not protein stability, position Ser140 plays an important role as its auto-phosphorylation triggers dissociation of the dimer after CHK2 activation [48]. Phosphomimetic mutations S140D and S140E and phospho-dead mutation S140A exhibited damaging map scores of −1.1, −0.6, and 0.2, respectively. It has been suggested that the wild-type *CHEK2* allele can allow activation and dissociation of an S140-mutated CHK2 partner [48], potentially explaining how we might observe a functional impact in our (haploid) setting but limited impact on protein stability. Taken together, these results support the previously-suggested importance of CHK2 dimerization [9] and that our map provides information about residue function beyond impacts on protein stability.

### Inferring residues with roles in CHK2 catalytic activity

Several well-established motifs–including GxGxxG, VAIK, HRD, DFG, and APE– coordinate kinase activation by influencing CHK2 conformation and binding of the substrate and cofactors ATP and magnesium [34] [37]. During kinase activation, after phosphorylation of Thr68 by ATM and subsequent CHK2 homodimerization, the auto-phosphorylation of residues Thr383 and Thr387 in the activation T-loop (positions 371-391) affects several residues corresponding to the above kinase motifs. For example, phosphorylation of Thr383 and Thr387 would be expected to promote electrostatic interaction with the positively charged catalytic HRD residues (345-347) and accompanying conformational change of the DFG residues (368-370) [49]. Switching from the inactive ‘DFG-out’ conformation to an active ‘DFG-in’ state helps to create the nucleation site and enable binding of ATP and magnesium [50]. Given the importance of these residues, damaging scores for HRD and DFG were expected, however, several flanking residues surrounding both HRD (343 to 344 and 348 to 355) and DFG (364 to 367 and 371 to 373) also showed damaging scores, suggesting the possibility that these residues may also be important for kinase activation. Future structural analysis, exploring electrostatic dynamics of the HRD motif as well as the fraction of time spent in the ‘DFG-in’ versus ‘DFG-out’ conformations for different *CHEK2* variants, could elucidate residues necessary for catalysis.

Interestingly, the T-loop itself showed that substitutions at some (but not all) positions impacted protein function. The activation T-loop (positions 371-391) is characterised by two amphipathic alpha-helices spanning positions 377-386 and 392-402 [34]. Many positions in the T-loop (including 374 to 380, 384, 388, 389, & 391) displayed surprising tolerance to most substitutions. Indeed, both of the involved alpha-helices had a segment displaying tolerance to a range of mutations, including helix-breaking prolines. Discovering tolerated positions at the C-terminal end of the activation T-loop was somewhat surprising considering that the conserved APE motif is directly downstream at positions 392 to 394, and that changing the size and charge of residues adjacent to APE might be expected to impact its ability to stabilise the kinase C-lobe during activation. Indeed, a role in stabilising the C-lobe may explain why residues in an alpha helix C-terminal to APE (405-423) were intolerant to substitution. In any case, our results show that many residues in the functionally important and generally-conserved T-loop are tolerant to substitutions.

Conformational changes in CHK2 following T-loop phosphorylation affect the GxGxxG motif (positions 227 to 232) in the glycine rich loop [34]. In the active CHK2 conformation, the GxGxxG motif acts as a flexible clamp, anchoring and orienting ATP for transfer of its phosphate group [51]. Interestingly, residues 220 to 226 preceding the GxGxxG motif were sensitive to polar substitutions (median score 0.0) but fairly tolerant to hydrophobic variants (median 0.8). Positions C-terminal to the GxGxxG motif may also play a role, based on damaging scores for most substitutions at positions 234, 237 and 238. Substitutions with bulky side-chains may reduce flexibility of the glycine rich loop and limit its ability to anchor ATP.

Conformational changes in CHK2 following T-loop phosphorylation also enable the VAIK (positions 246 to 249) salt bridge [52] between Lys249 and Glu273 across an intervening disordered loop from position 255 to 268. This stabilises the glycine rich loop and thereby enables ATP binding [34]. As expected, the disordered loop is relatively tolerant to variation (median score 0.7). Positions adjacent to the VAIK salt bridge including 239 to 245 and 250 to 252, were surprisingly tolerant to substitutions (median score 0.7).

Further downstream from the VAIK motif, positions 253 to 258 were somewhat intolerant to variation and appeared to match an enrichment of arginine and lysine residues typical of Protein Kinase C (PKC) substrates. PKC is known to regulate proliferation [53] and DNA damage repair [54]. However, the nearby Ser260 phosphorylation site is not conserved and, with a median map score of 1.2, appears to tolerate substitutions. Therefore, damaging scores at positions 253 to 258 more likely suggest roles for the positions in VAIK salt bridge formation and ATP binding. Together, our map sheds light on additional positions that may be important for functions of the glycine rich loop and VAIK salt bridge that work together to stabilise ATP for phosphoryl transfer during catalysis.

Several amino acids in CHK2 have been shown to coordinate ATP binding more directly. Crystal structures of CHK2 complexed with ADP or the ATP analogue DBQ identified hydrogen bonds involved in ATP binding at positions Lys249, Glu302, Met304, Glu308, Glu351, Asn352, and Asp368 [34]. Our map showed that most of these residues are intolerant to substitutions. The exceptions (Glu302 and Met304 which had median scores of 1.2 and 0.7 respectively) are understandable given that Glu302 and Met304 form hydrogen bonds via backbone rather than side-chain atoms. Adjacent substitutions (at positions 301, 306, 307, and 309) also exhibited damaging scores, suggesting that these residues may indirectly promote hydrogen bonding by Glu302, Met304 and Glu308. More specifically, substituting small and uncharged residues for bulky amino acids at these positions may affect CHK2 function by steric hindrance, occluding the hydrogen binding sites needed for ATP binding. For the remaining ATP binding residues (Lys249, Glu351, Asn352, and Asp368) distinguishing between their roles in catalysis and ATP binding is challenging, given that Lys249, Asn352 and Asp368 overlap with the VAIK, HRD+5, and DFG mutational hotspots respectively.

### Potential clinical applications for the CHK2 variant effect map

While loss of CHK2 function has been linked to several cancer types including prostate and colorectal, we focus here on the association with breast cancer. In contrast with the predisposition to breast cancer that is well-established for truncating CHK2 variants, the risk associated with missense variants is less often clear [55]. Some CHK2 missense variants, such as I157T and S428F (which received map scores of 0.3 and 1.2 respectively) have been reported to convey a modest (<1.5 fold) elevation of breast cancer risk. By contrast, R117G, which reportedly conveys a >2-fold breast cancer risk that is on par with truncating variants [56], showed a correspondingly low map score of - 1.0. Indeed, a recent analysis showed that *CHEK2* variants found to be dysfunctional in mammalian cell-based assays (found in 0.5% of breast cancer patients) were associated with an increased risk of breast cancer, while functionally normal or mildly dysfunctional variants (found in 2.2% of patients) were not associated with a clinically relevant increased risk of cancer [57]. Thus, a CHK2 variant effect map could help stratify variants by clinical risk and thereby focus management of clinical resources.

Improved variant interpretation and risk estimation can enable personalised medicine, with surveillance and treatment plans that depend on an individual’s genotype. For example, heightened screening may be warranted for patients with pathogenic germline *CHEK2* variants. It has been estimated that establishing regular MRI and mammograms based on *CHEK2* genotype could reduce breast cancer mortality by over 50% [58]. Given the suggestion from a recent phase II clinical trial that *CHEK2* breast cancers respond less well to poly-adenosine diphosphate ribose polymerase inhibitors, there is the future potential for knowledge of *CHEK2* genotype to inform therapy [59]. Patients with high-risk CHK2 variants and a strong family history of breast cancer may benefit from a pre-emptive or contralateral risk-reducing mastectomy [55]. Together, these results support the potential clinical value of our proactive CHK2 variant effect map.

### Limitations/caveats of the map

An important caveat of our map is that it may not detect the impact of variants on human CHK2 functions that are not carried out by the yeast Rad53 ortholog or which are not enabled for human CHK2 in yeast. For example, CHK2 may have (and depend on) post-translational modifying activities in human cells that are not present in yeast under our assay conditions. Kleiblova *et al.*[38] hypothesised that post-translational modifications can influence CHK2 catalytic activity in human cells. Using a mammalian cell-based *in vivo* assay they demonstrated that, when CHK2 undergoes physiological post-translational modifications, the protein has a greater ability to phosphorylate KAP1-S473 compared to unmodified recombinant CHK2 tested in an in-vitro assay. Post-translational modifications like phosphorylation and ubiquitination are both important for CHK2 function as a DNA damage checkpoint [60]. Both CHK2 and its yeast homolog, Rad53, are phosphorylated after translation but only CHK2 is ubiquitinated by E3 ligases, which serve to regulate CHK2 protein stability [61] [62]. There are also many differences in the downstream pathways between human CHK2 and yeast Rad53. Many CHK2 substrates, including Cdc25A/B/C, Pik3 kinase, E2F1 transcription factor, BRCA1/2, and p53, do not have clear yeast counterparts [5,63]. Another limitation is that our yeast assay was conducted at a temperature suitable for optimal yeast growth which is 30°C instead of 37°C, so that thermodynamic stability of human CHK2 variants expressed in the yeast model may differ from human physiological conditions [64]. This could explain why some CHK2 variants exhibiting intermediate functional effects in a mammalian system [9] appear functional in a yeast-based assay [11].

In our assay, human *CHEK2* cDNA was expressed under the constitutive *ADH1* promoter. Although this promoter is often considered to have moderate strength, we cannot be sure that this does not represent overexpression of the protein, such that some variants that would be mildly dysfunctional at physiological expression levels in a human cell could provide completely sufficient total activity when over-expressed in yeast. Conversely, some variants might be toxic to yeast when overexpressed but tolerated at physiological expression levels in human cells.

A further limitation of any cDNA rescue assay is that it will miss some purely non-coding effects of CHK2 coding variants, e.g. on splicing efficiency. Also, observations that a nonsense codon is tolerated in our assay should not be taken as strong evidence that the variant will be tolerated in humans, given that strength of nonsense-mediated decay effects can depend on the presence of downstream introns.

Despite all of these caveats, assays based on the expression of human cDNA in yeast can provide excellent empirical performance in identifying pathogenic variation [22] [19] [32] [65]. Here, we showed that approximately half of CHK2’s known pathogenic missense variants could be identified at a stringency achieving 90% balanced precision, i.e., if a balanced training set is applied, we would expect that 90% of the map’s inferred-pathogenic variants would in fact be pathogenic.

## Conclusion

This study provides the largest resource to date of *in vitro*-based functional assays of *CHEK2* missense variants, enabling both biochemical insights and representing a proactive assessment of nearly all possible missense variants of *CHEK2*, with potential to enable more rapid and accurate clinical action of *CHEK2* clinical variants.

## Materials and methods

### Yeast strains and plasmids

A *Saccharomyces cerevisiae* strain (*MAT*a *sml1*Δ::*kanR rad53*Δ::*hygR*) was used as a host for the *CHEK2* variant library. The double deletion strain was generated by PCR replacement of the *RAD53* gene with the Hygromycin selectable marker (cassette) in the single deletion strain (*MAT*a *sml1*Δ::*kanR*) derived from the yeast knockout collection [66].

The *CHEK2* open reading frame (ORF) clone, corresponding to UniprotKB accession O96017, was obtained from the Human ORFeome v8.1 library [67]. A Gateway compatible yeast expression vector, pHYC-Dest2 (CEN/ARS-based, *ADH1* promoter, and *LEU2* marker), was used for the complementation assay.

Wild-type reference or mutated disease-associated versions of the *CHEK2* ORFs were transferred into pHYC-Dest2 by Gateway LR reactions. After confirmation of ORF identity and expected mutations by Sanger sequencing, the expression clones were transformed into the double deletion yeast strain in parallel with an ‘empty’ expression vector control (bearing the counterselectable ccdB marker controlled by a bacterial promoter).

### CHEK2 Yeast complementation assay

Single colonies of yeast transformed with vectors expressing *CHEK2* cDNAs were picked from SC-Leu+Kan+Hyg with 2% glucose plates and grown at 30°C to saturation (overnight) in liquid media SC-Leu+Kan+Hyg with 2% glucose. Each culture was then adjusted to an OD600 of 0.2 and diluted 1:5 with 5 serial dilutions. 4µl of these cultures were spotted on agar plates of SC-Leu containing 2% glucose and a final concentration of 0.007% MMS (spotting assay). The plates were incubated at 30°C and after imaging, the comparison of the effect of MMS on growth/fitness of yeast–carrying either *CHEK2* variants, *CHEK2*-WT, or an empty vector–was made starting from day 3 (Fig.1).

### Construction of codon randomised CHEK2 variant libraries

As a first step of the TileSeq framework, a pooled random-codon mutagenesis method (<u>P</u>recision <u>O</u>ligo-<u>P</u>ool based <u>Code</u> Alteration or POPCode) [13] was used to construct the *CHEK2* variant libraries. Mutagenesis was separately applied in four regions of the *CHEK2* ORF (which has a total length of 1632 bp) that are each approximately 150 amino acids long (Supplementary Figure [Regions and Tiles]), following each step of the framework below separately to generate four distinct regionally-mutagenized libraries. Oligonucleotides (28-38 bp long), with centered NNK-degenerate codons that cover the entire *CHEK2* coding sequence, were designed based on optimal melting temperatures using the POPcode oligo suite tool [13]. Oligonucleotides for each of the four *CHEK2* regions were pooled and phosphorylated to perform regional POPCode mutagenesis. Phosphorylated oligonucleotides were annealed to a uracilated full-length template of wild-type *CHEK2* using KAPA HiFi Uracil+ DNA polymerase (KapaBiosystems) and a mix of dNTP/dUTP from each regional oligo pool. After annealing, KAPA HiFi Uracil+ DNA polymerase (KapaBiosystems) was used to fill in the gaps and Taq DNA ligase (NEB) was applied to seal the nicks. The samples were treated with Uracil-DNA-Glycosylase (UDG) from NEB to degrade the original uracilated template. The newly POPCode-mutagenized strand was then amplified with primers containing attB sites. The mutagenized attB-PCR products were next cloned *en masse* into the entry vector pDONR223 by Gateway BP reactions, generating regional Entry libraries that had ∼150,000 clones per region, with the goal of achieving high complexity, i.e., so that each variant would be present in an average of 50 or more independent clones. These Gateway-entry clone libraries were then transferred to a pHYC-Dest2 expression vector by *en masse* Gateway LR reactions, generating regional expression libraries from >300,000 transformants per region in order to maintain pool complexity. Both Gateway-Entry and -expression library transformations used NEB5α *E. coli* cells (NEB), with selection on LB agar plates using spectinomycin and ampicillin, respectively. Both Gateway-Entry and -expression libraries and the yeast transformants were grown on 245×245mm^2 bioassay dishes (Corning). Finally, regional expression libraries were transformed into the *S. cerevisiae* double mutant strain *sml1*Δ *rad53*Δ using the EZ Kit Yeast Transformation kit (Zymo Research) to obtain ∼1,000,000 clones per region, thus maintaining library complexity. Yeast transformants for each regional library were pooled separately and grown in Synthetic Complete (SC) without Leucine (USBiological) as the non-selective media. In an initial attempt at mutagenesis of the fourth region, frequent frameshift mutations were observed (data not shown). After attributing this to a TG-rich segment in the last 23 nucleotides of the CHEK2 ORF, we designed a synthetic clone that was codon-optimised to limit potential secondary structures before repeating regional POPcode mutagenesis for the fourth region. No enrichment for frameshifts was subsequently observed in the final version of the fourth regional mutagenesis library.

### Multiplexed assay for CHK2 variant function

The yeast-based functional complementation assay we used to evaluate *CHEK2* missense variants similar to that of Roeb *et al.,* 2012, who assayed recovery of cells from DNA damage treatment (0.014% MMS in liquid media). For this study, we selected cells using SC-Leu solid-agar media plates with 0.007% MMS. More specifically, from the yeast transformants grown in non-selective conditions, two replicates of ∼9M cells from each of the regional transformant pools were plated onto selective SC-Leu media-plates with an MMS concentration of 0.007% and incubated at 30°C for 3 days. After pooling the colonies of each replicate, plasmid DNA was extracted from ∼4M cells per region from both non-selected and selected pools and used for subsequent PCR amplification of specific targeted tiles. Controls were assayed, in parallel, using the *S. cerevisiae sml1Δrad53Δ* strain transformed with an empty pHYC-Dest2 expression vector, as the negative control, and wild-type *CHEK2* in the pHYC-Dest2 vector, as the positive control. The control strains were grown in petri dishes of solid non-selective and selective media during the same incubation period as the large-scale variant library regional assays.

### Quantifying variant abundance

Each region was conceptually divided into four tiles, each approximately 151 nucleotides long. Each tile was amplified from the plasmid DNA extracted from each pool of yeast from non-selective and selective conditions, amplifying targeted tiles with primers carrying an Illumina sequencing-adaptor binding site (Table S4). Next, a unique Illumina sequencing adapter was added to each tile via a subsequent “indexing PCR”. Equal amounts of the tiled indexed PCR products were pooled together and the resulting pooled library of an expected size of ∼300bp was purified using a 4%EX e-gel (Life Technologies) followed by MinElute Gel Extraction (Qiagen). After the library quantification via NEBNext Library Quant Kit (NEB), paired-end sequencing was performed on the tiles of each region with a sequencing depth of ∼2 million reads per tile using an Illumina NextSeq 500 instrument via a NextSeq 500/550 High Output Kit v2.

### Deriving functional scores for variants

An analysis pipeline, called TileseqMave (version 1.0.0), was used for the generation of the variant effect map (code, installation instructions and documentation are available on Github at https://github.com/rothlab/tileseqMave. TileseqMave uses bowtie2 [68] forward and reverse reads to the *CHEK2* reference sequence. For each divergent base-call, the posterior probability of being a true variant is calculated. Variants with posteriors exceeding a threshold of 0.9 are counted. For each codon change, a "marginal count" is calculated, that is the number of times the change was observed irrespective of other co-occurring variants. To calculate the frequency of each codon change, the marginal count for each mutation is normalised by its "effective sequencing depth", i.e. the number of reads in which the variant call at the given position was decidable. Then, an error-corrected enrichment log-ratio is calculated by subtracting the frequency of the variant in the corresponding WT control from the post- and pre-selection frequencies and then calculating the log ratio between the latter. (Where multiple sequencing runs were required to obtain sufficient read counts, we subtracted counts from matched WT control libraries before aggregating counts.) Finally, the enrichment log-ratio is rescaled such that, after rescaling, synonymous variants have a median score of 1 and nonsense variants a median score of 0. For each variant, measurement error is regularised using the method described by Baldi and Long [69] and propagated via bootstrapping.

To include only well-measured results, we filtered out variants that were seen in less than 10 reads and whose pre-selection frequencies were statistically indistinguishable from those in the WT control. We also removed variants for which replicates diverged by more than three times the amount expected based on their Poisson variance given their underlying read count.

As an optional addition in the supplemental tables and figures, we performed imputation and refinement of the baseline original scores. Scores for all possible amino acid substitutions were imputed based on the median of available scores from the five most similar amino acid substitutions (according to BLOSUM100). Imputed values were then refined by combining imputed and original values, using a weighted average with weights that maximised correlation of resulting imputed/refined scores with the variant effect predictor VARITY_R. Further filtering to isolate high-quality variants was performed for the clinical analysis by removing variants where the confidence interval of a variant’s score plus and minus error overlapped with the midpoint between synonymous and nonsense (i.e. a score of 0.5).

### Conservation, Solvent Accessibility, and Protein Stability

To quantify evolutionary conservation across *CHEK2*, we used ConSurf [70] [71]. As ConSurf considers structural homology when choosing homologs for comparison, we provided a predicted structure [using AlphaFold version 2022-11-01, Monomer v2.0 pipeline; UniProt O96017]. Consurf assigned a conservation score to each position along *CHEK2* ranging from highly conserved scores below −0.47 to lowly conserved scores above 0.91. The median functional score for every position with a conserved grade was compared to functional scores in non-conserved positions using a Wilcoxon rank-sum test.

Solvent accessibility was calculated using FreeSASA [version 2.02 October 22nd, 2017] using the above-mentioned CHK2 structure [72]. Positions with relative side-chain accessibility to solvent below 20% were considered inaccessible and positions greater than 50% accessible. Median functional scores for positions in the FHA domain (residues 92-205) and the kinase domain (residues 212-501) were stratified into solvent-accessible and -inaccessible positions then plotted and compared by Wilcoxon rank-sum test.

ΔΔG thermostability predictions were made for all possible CHK2 missense variants using the ddGUN software [version 0.0.2] [73]. ΔΔG scores below −1 destabilise the protein while scores near 0 retain structure. With the median functional score and median stability score at each position as the input, we performed a moving window analysis across CHK2 that took the median value from a bin width of five positions and moved at a step-size of one residue. Correlation between functional score windows and ΔΔG stability scores was evaluated by PCC.

### Reference Set of Clinically Annotated Variants

A set of clinically-annotated *CHEK2* variants was provided by Invitae, using their Sherloc v6.0 variant classification system which captures evidence of strength in terms of points towards pathogenicity or benignity [41]. Variants with at least four pathogenic points were included in the positive reference set (including variants annotated as either pathogenic and likely pathogenic variants) and variants with at least three benign points were included in the negative reference set ( including variants annotated as either benign or likely benign). From our sets of 21 positive and 39 negative reference variants, 12 positive and 28 negative variants had high-confidence functional scores.

### Calibrating Log-Likelihood Ratios to ACMG Evidence Strength

To derive estimates of the strength of evidence towards pathogenicity annotation that are both compatible with a Bayesian clinical annotation approach and are more continuously quantitative than points-based approaches, we estimated the log-likelihood ratio of pathogenicity (LLR) for each variant. More specifically, probability density functions were separately estimated for scores from the positive and negative reference variant sets using kernel density estimation (using a gaussian kernel with a bandwidth determined by biased cross-validation). Then, for each variant, the LLR could be calculated as the log-ratio of the two probability densities regularised against a uniform distribution. LLR values were then compared to threshold values defined by the ACMG/AMP variant classification framework using the strategy of Tavtigian *et al.* [43] as subsequently adapted [32].

### Structural modelling and analysis

We used OpenPyMol to visualise structural models of CHK2 using previously reported structural models (PDB:3i6u [33], PDB:2CN5 [34], and PDB:2CN8 [34]. Structural models were coloured according to the median variant effect map score for each residue.

## Supporting information

Table S5. All CHEK2 functional scores

## Acknowledgements

The authors gratefully acknowledge funding from the National Human Genome Research Institute of the National Institutes of Health (NIH/NHGRI) Center of Excellence in Genomic Science (CEGS) Initiative (HG004233 and HG010461), the NIH/NHGRI Impact of Genomic Variation on Function (IGVF) Initiative (UM1HG011989), the NIH/NHLBI grant (HL164675), the Canada Excellence Research Chairs (CERC) Program, and a Canadian Institutes of Health Research Foundation Grant to F.R.

## Supplementary material

**Table S1.** Fractions of amino acid substitutions that were sufficiently well represented in the “non-select” expression library (after transformation into yeast but not subjected to selection) to be considered well-measured. Here, we considered amino-acid substitutions to be well measured if at least 50 read counts were observed in the non-select library. Fraction of all possible codon changes considers missense and nonsense variants across all *CHEK2* positions.

| Region | Amino Acid (AA)<br>Changes Per Clone | Fraction of all possible AA<br>changes that are<br>well measured | Fraction of all possible 1<br>nt-accessible AA<br>changes that are well<br>measured |
| --- | --- | --- | --- |
| 1 | 0.51 | 86% | 99% |
| 2 | 0.17 | 81% | 100% |
| 3 | 0.37 | 92% | 100% |
| 4 | 0.22 | 62% | 98% |

**Table S2.** List of residues that had been previously identified as important for CHK2 activation on the basis of CHK2 crystal structures, based on proximity to either complexed ADP or the competitive ADP inhibitor debromohymenialdisine (DBQ). Columns of median and mean score and the number of variants above or below a score of 0.5 consider all well-measured missense variants at each position.

| **CHK2 amino acid position** | **Nucleotide and inhibitor binding sites** | **Median** | **Mean** | **Number of variants with score > 0.5** | **Number of variants with score < 0.5** |
| --- | --- | --- | --- | --- | --- |
| **G227** | ADP | 0.19 | 0.49 | 8 | 9 |
| **S228** | ADP | 0.50 | 0.49 | 8 | 8 |
| **G229** | ADP | 0.06 | 0.33 | 4 | 9 |
| **G232** | ADP | -0.35 | -0.38 | 0 | 16 |
| **V234** | ADP and DBQ | -0.39 | -0.13 | 4 | 13 |
| **A247** | ADP and DBQ | -0.26 | -0.26 | 2 | 14 |
| **K249** | ADP and DBQ | -0.31 | 0.08 | 3 | 12 |
| **L301** | DBQ | 0.23 | 0.32 | 6 | 11 |
| **E302** | ADP and DBQ | 1.12 | 1.23 | 18 | 0 |
| **L303** | ADP | 1.20 | 1.37 | 15 | 0 |
| **M304** | ADP and DBQ | 0.70 | 0.65 | 10 | 7 |
| **E308** | ADP and DBQ | 0.04 | -0.11 | 3 | 14 |
| **E351** | ADP and DBQ | 0.22 | 0.07 | 4 | 12 |
| **N352** | ADP and DBQ | 0.04 | -0.27 | 0 | 18 |
| **L354** | ADP and DBQ | -0.22 | 0.17 | 6 | 12 |
| **D368** | ADP | -0.62 | -0.38 | 1 | 16 |

**Table S3.** Mutational Hotspots. Mutational hotspots identified in *Hudson et al. 2018* [37] were separated according to their correspondence to distinct kinase activation motifs. For each hotspot, the table indicates the amino acid position relative to the beginning (negative values) or end (positive values) of the nearest motif, as well as the median and mean of original functional scores at these positions. Of the 23 CHK2 hotspots identified, all 16 that fell within a named motif (APE, VAIK, HRD, DFG and the three G positions within the GxGxxG motif) scored as intolerant to variation in our map (i.e., with median scores below 0.5).

| Hotspot | CHK2<br>amino acid<br>position | Median | Mean | Number of<br>variants<br>with score<br>> 0.5 | Number of<br>variants<br>with score<br>< 0.5 |
| --- | --- | --- | --- | --- | --- |
| APE | 392 | -0.06 | -0.02 | 5 | 14 |
|  | 393 | -0.76 | -0.56 | 2 | 13 |
|  | 394 | -0.34 | -0.28 | 2 | 12 |
| Salt bridge E | 273 | 0.43 | 0.41 | 1 | 2 |
| VAIK | 246 | 0.35 | 0.53 | 9 | 10 |
|  | 247 | -0.26 | -0.26 | 2 | 14 |
|  | 248 | -0.05 | 0.25 | 8 | 10 |
|  | 249 | -0.31 | 0.08 | 3 | 12 |
| HRD | 345 | -0.11 | -0.27 | 2 | 14 |
|  | 346 | 0.05 | -0.04 | 2 | 15 |
|  | 347 | -0.55 | -0.45 | 2 | 15 |
| GxGxxG | 227 | 0.19 | 0.49 | 8 | 9 |
|  | 228 | 0.50 | 0.49 | 8 | 8 |
|  | 229 | 0.06 | 0.33 | 4 | 9 |
|  | 230 | 1.27 | 1.12 | 3 | 1 |
|  | 231 | 1.12 | 0.98 | 13 | 5 |
|  | 232 | -0.35 | -0.38 | 0 | 16 |
| DFG | 368 | -0.62 | -0.38 | 1 | 16 |
|  | 369 | -0.42 | -0.57 | 0 | 18 |
|  | 370 | 0.07 | 0.07 | 2 | 6 |
| HRD + 5 | 352 (Asn) | 0.04 | -0.27 | 0 | 18 |
| APE – 6 | 386 (Gly) | 0.05 | -0.03 | 5 | 14 |
| HRD – 6 | 339 (His) | 0.86 | 0.72 | 10 | 7 |
| **HRD + 7** | 354 (Leu) | -0.22 | 0.17 | 6 | 12 |
| **HRD – 7** | 338 (Leu) | 0.56 | 0.49 | 7 | 7 |
| **APE – 3** | 389 (Thr) | 1.10 | 1.13 | 15 | 4 |

**Table S4.**
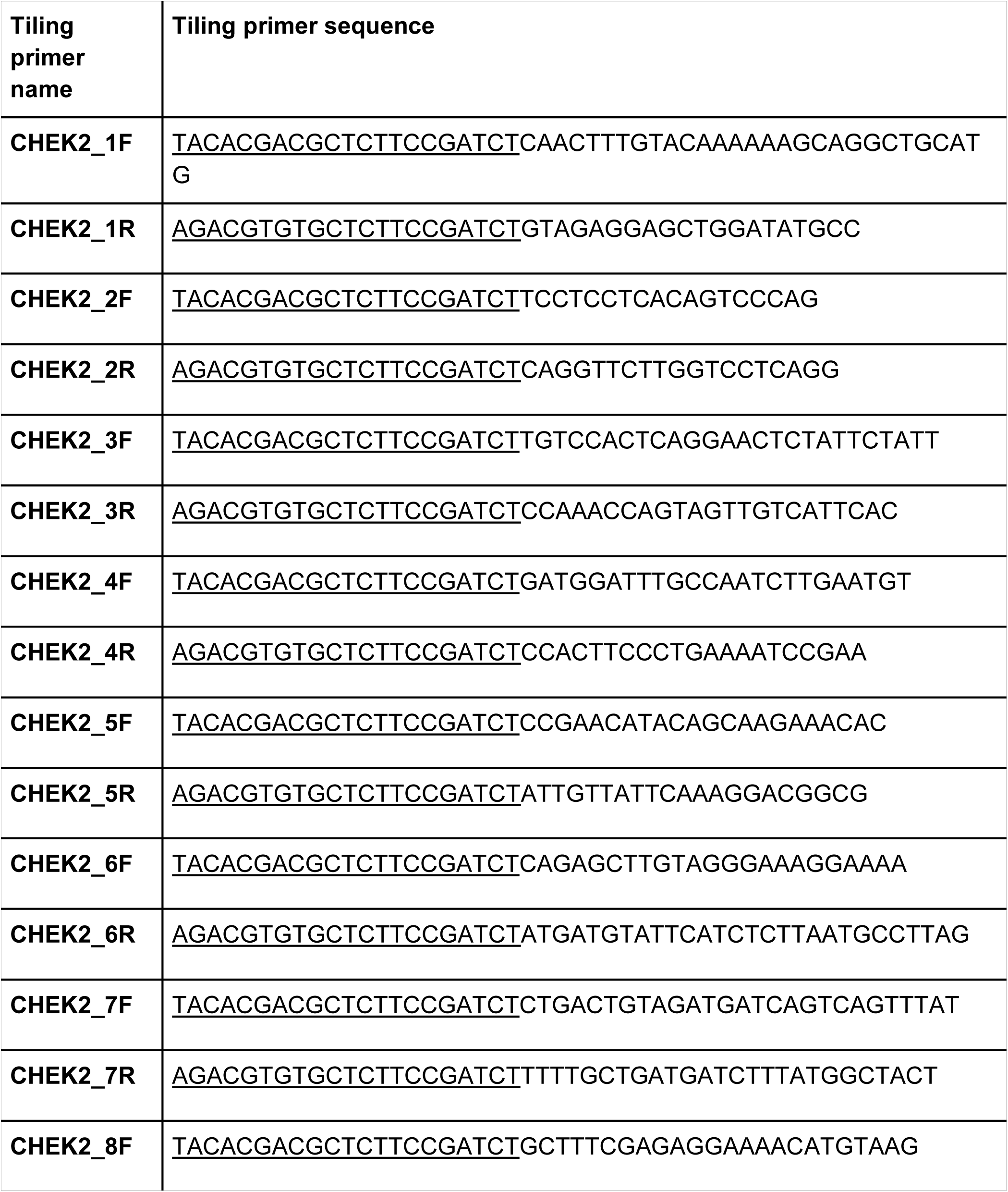
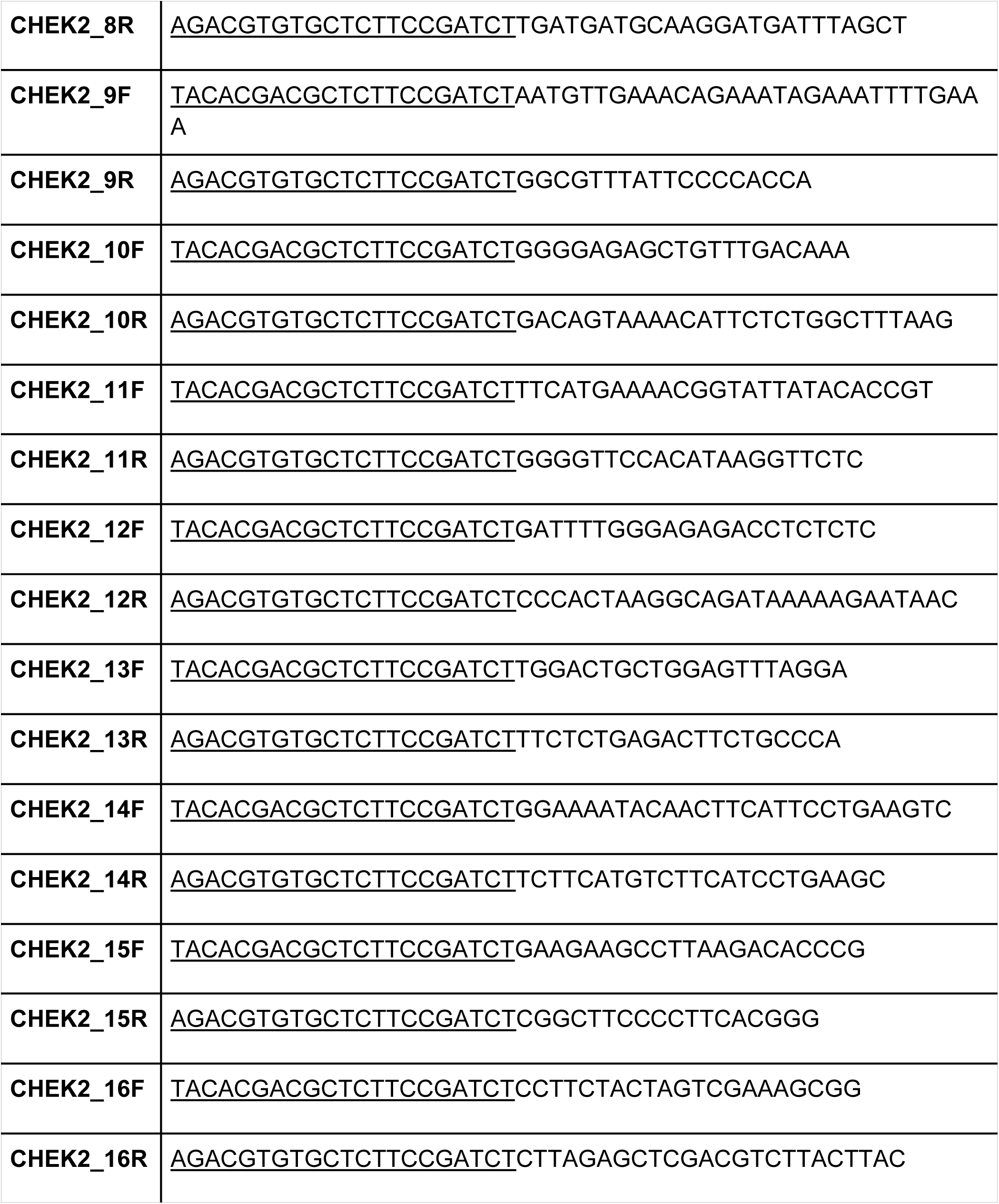
Tiling primers. For each of the plasmid libraries from non-selective and selective pools, primers carrying a binding site for Illumina sequencing adaptors ( underlined in the sequences in the table) were used to amplify short template amplicons (tiles) of ∼150 bp such that the union of tile positions internal to the priming sites covered the entire ORF.

## Supplementary Figures

**Figure S1.**
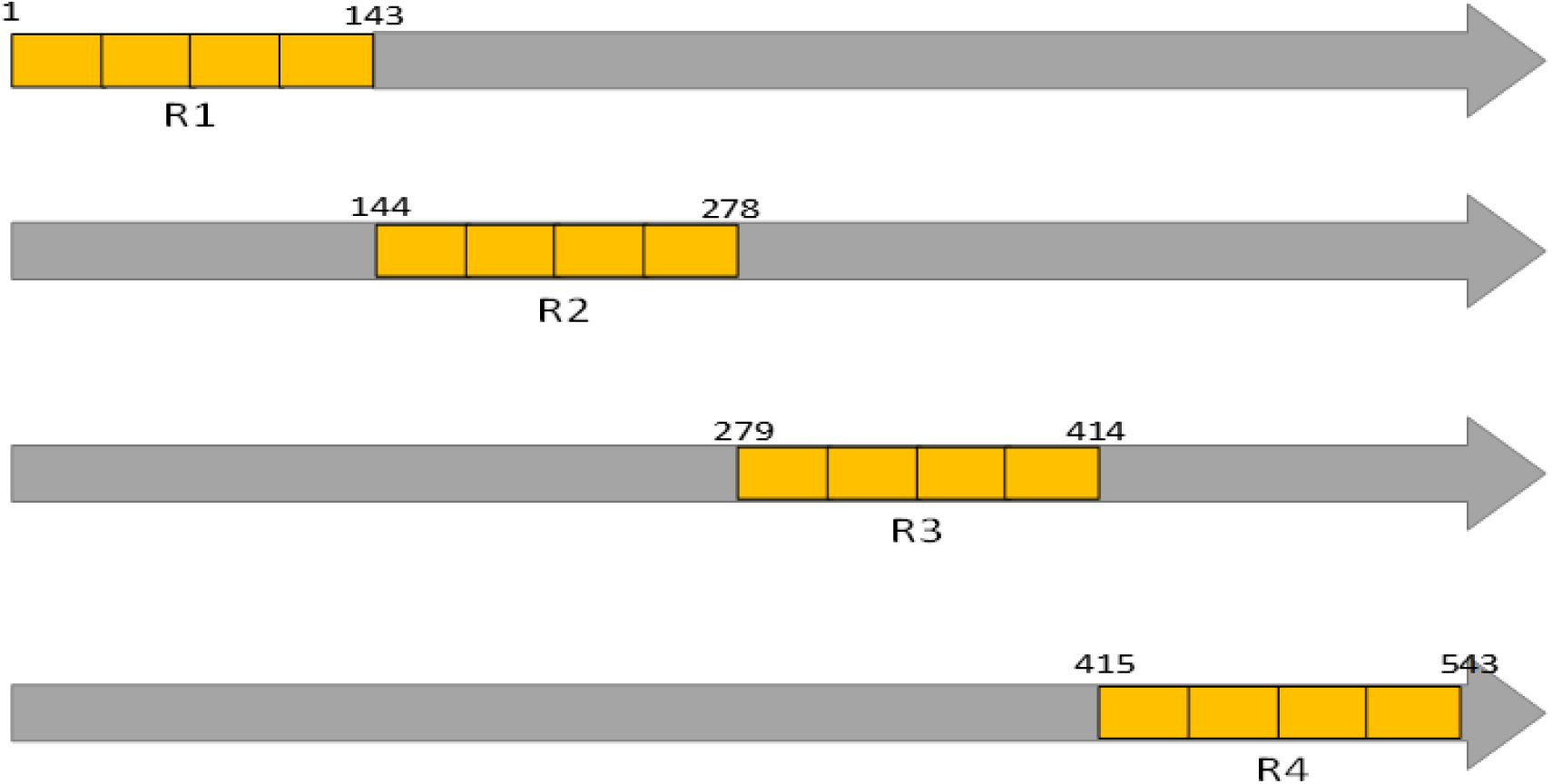
Mutagenized region of CHK2. We defined four regions of CHK2, corresponding to an average length of 150 AA each. Mutagenesis was targeted to each region in turn to generate four mutagenized libraries. For each region, we designed four sequencing tiles for the purpose of sequencing to estimate mutational frequencies before and after selection. The DMS-tileseq framework was followed separately for each regionally-mutagenized library.

**Figure S2.**
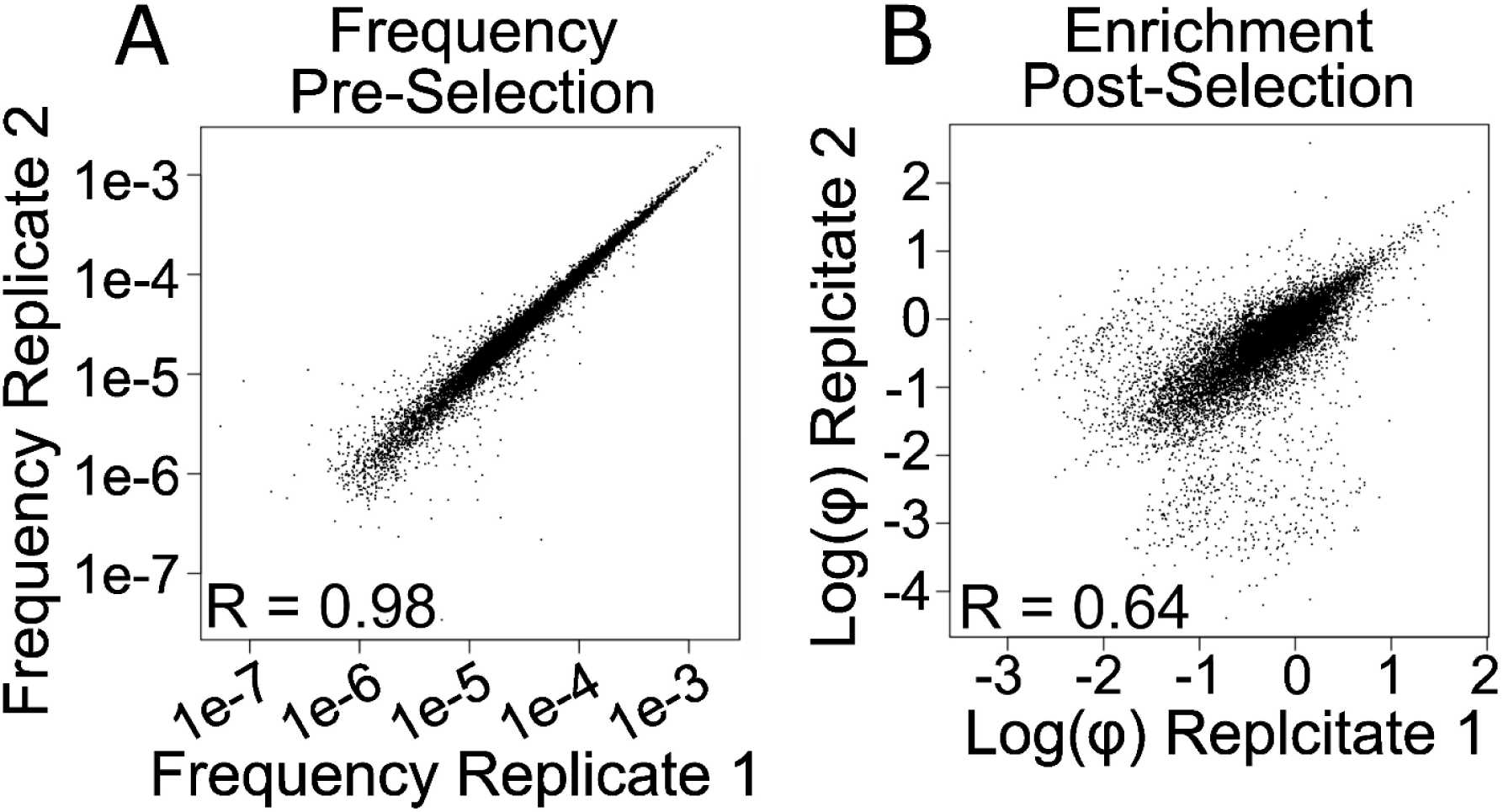
Replicate correlation across sequencing and selection. **A** To calculate the frequency of each *CHEK2* variant in the pool and estimate error rates associated with tiling PCR and sequencing, two independent sequencing libraries from the non-select condition (baseline media without MMS) were prepared (see “quantifying variant abundance” section in Methods). Correlation of variant frequency between replicate pools was assessed by Pearson correlation. **B** Two independent experimental replicates from the selective condition (media with 0.007%MMS) were performed and correlation of variant-specific log(φ) enrichment ratios across replicates (comparing the frequency of each variant in the non-select condition to the select condition) was assessed by Pearson correlation.

**Figure S3.**
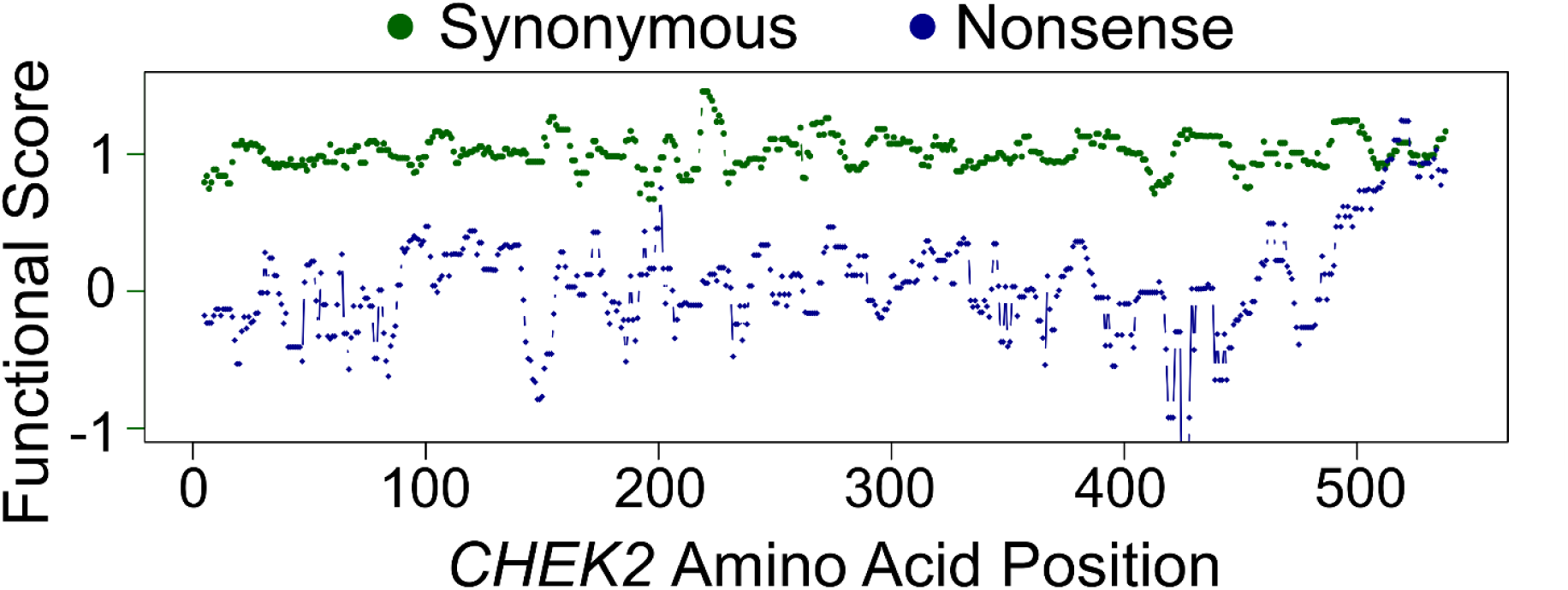
*CHEK2* MMS assay effectively separates synonymous and nonsense variants. A moving window analysis of synonymous and nonsense scores along CHK2 positions was performed to assess the experiment’s ability to separate neutral and loss-of-function variants. For each position evaluated, a window of 10 residues centered on that position was captured and median synonymous (green) and median nonsense (blue) scores were plotted. The majority of scores ranged from −1 to 1 with the exception of positions 425 to 427 for nonsense variants where scores dropped to between −2 to −3 (not shown).

**Figure S4.**
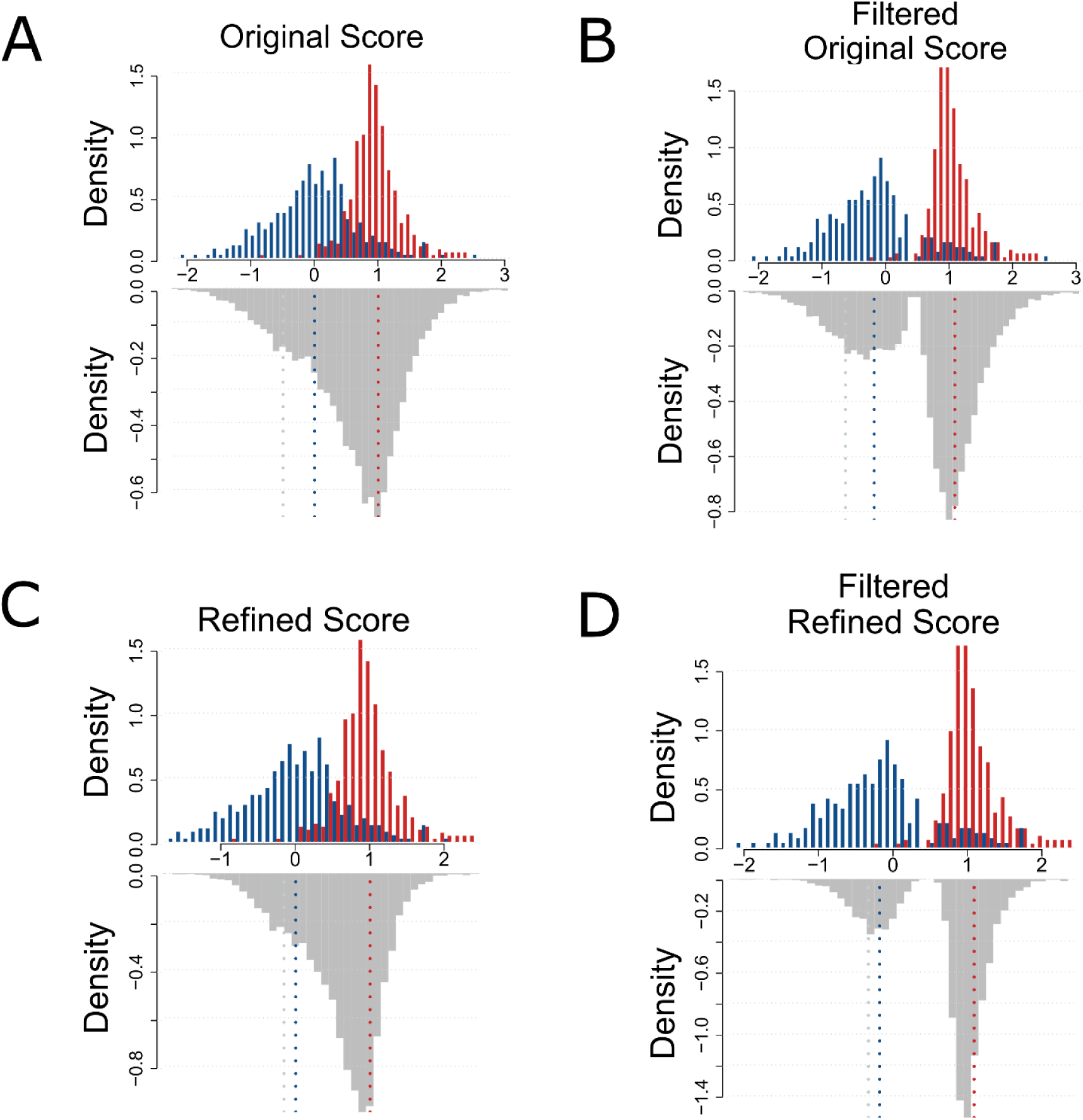
Distribution of *CHEK2* scores with and without refinement and filtering. **A** Functional scores for synonymous, nonsense, and missense variants from the original map were plotted as a histogram with scores on the x-axis and density on the y-axis. Synonymous variants are shown in red, nonsense in blue, and missense in grey. The median value for each variant type is shown as dotted vertical lines that match the colour code above. **B** Confidence interval filtering as described in the methods was applied to the original functional and plotted as described above. **C** Functional scores after imputation and refinement are also plotted as described above. **D** Imputed and refined functional scores underwent confidence interval filtering and plotted.

**Figure S5.**
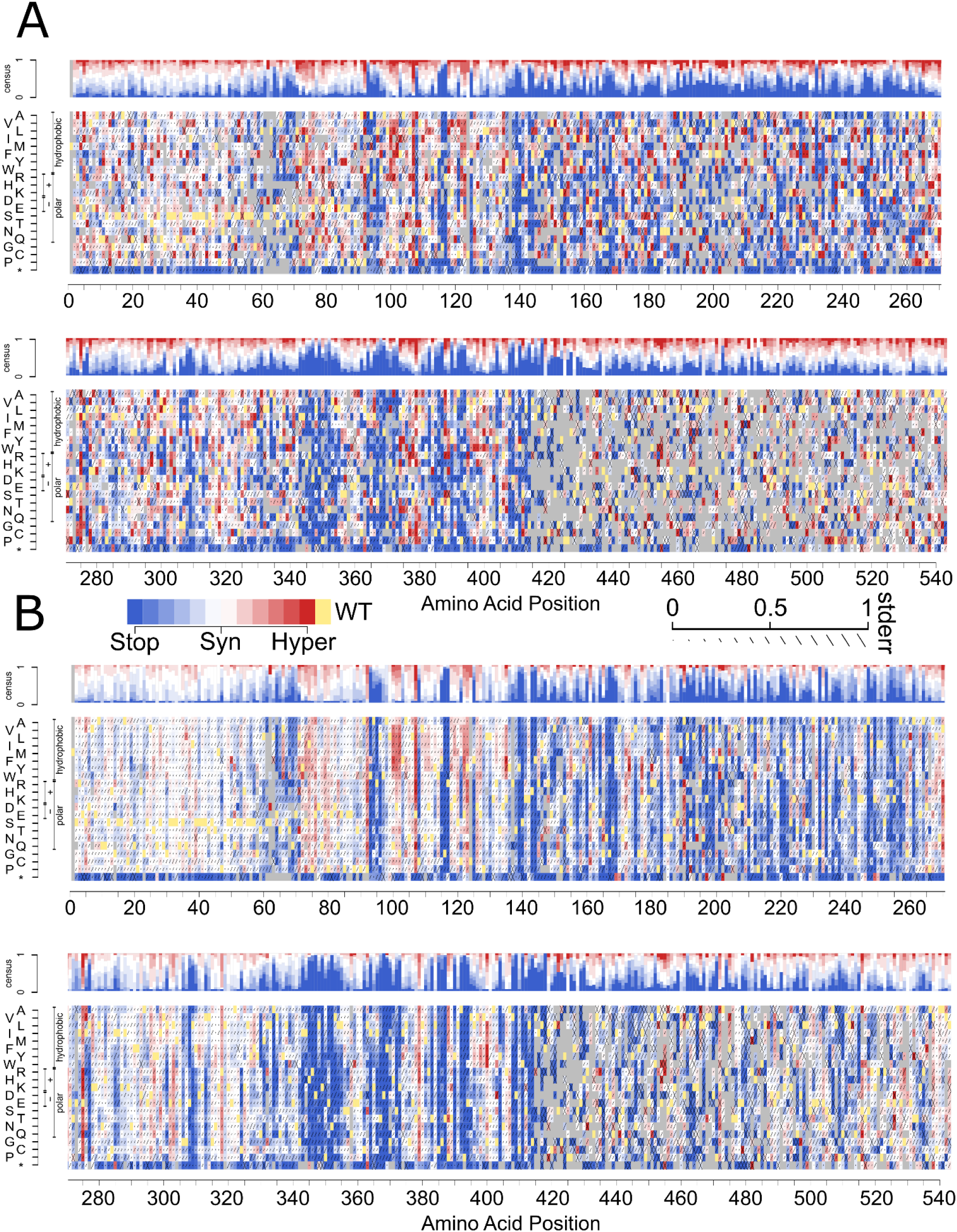
Full length original and refined CHK2 variant effect maps. **A** Functional scores derived directly from the original (unrefined) assays are shown as a heatmap with positions along CHK2 on the x-axis and amino acid substitutions on the y-axis. Blue indicates deleterious variants with scores near 0, white represents tolerated variants with scores near 1, red indicates apparently ‘hyper-complementing’ variants with scores above 1, yellow indicates the canonical wild-type amino acid, and grey indicates missing data. Within each cell, the estimated standard error is indicated by the total length of lines as described in the legend (e.g., 0 error is indicated with a dot, error of 1 is indicated by a full-length slash, and an error higher than 1 is indicated by an X. The census track along the top of the plot depicts the distribution of scores at each position. **B** Functional scores after imputation and refinement (see methods) are also plotted as described in panel A.

**Figure S6.**
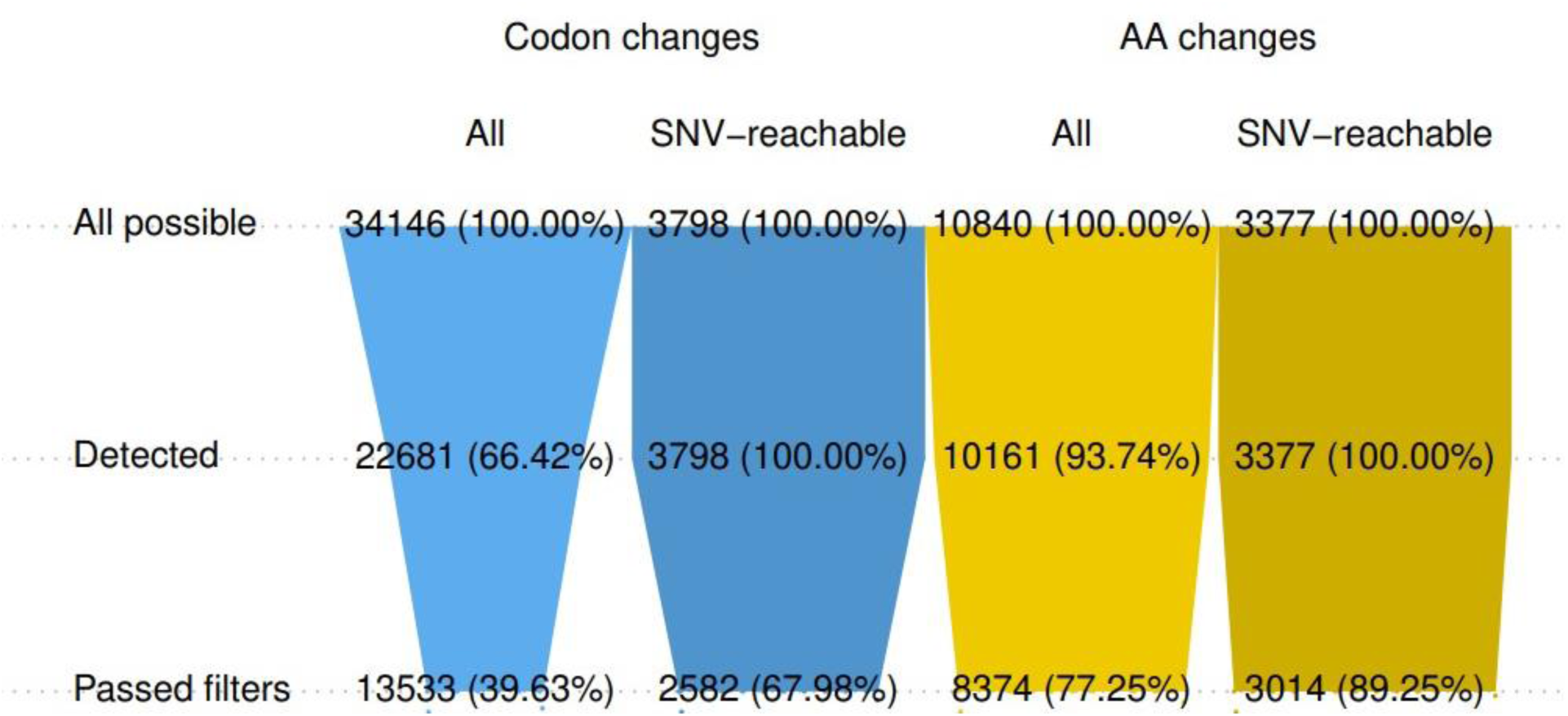
Coverage of all possible codons and amino acids in the CHK2 variant effect map. The proportion of variants in *CHEK2* covered by our experiment is indicated, either as: a fraction of all possible codon-level substitutions (lighter blue), a fraction of codon-level substitutions that can be reached via a single nucleotide change (darker blue); a fraction of all possible amino acid changes (lighter yellow); and a fraction of all possible amino acid changes that can be reached via a single-nucleotide change (darker yellow). “Detected” indicates variants that were observed during sequencing while “passed filters” refers to variants that were sufficiently well represented in the non-select library to be considered well-measured and were included in the final *CHEK2* variant effect map.

**Figure S7.**
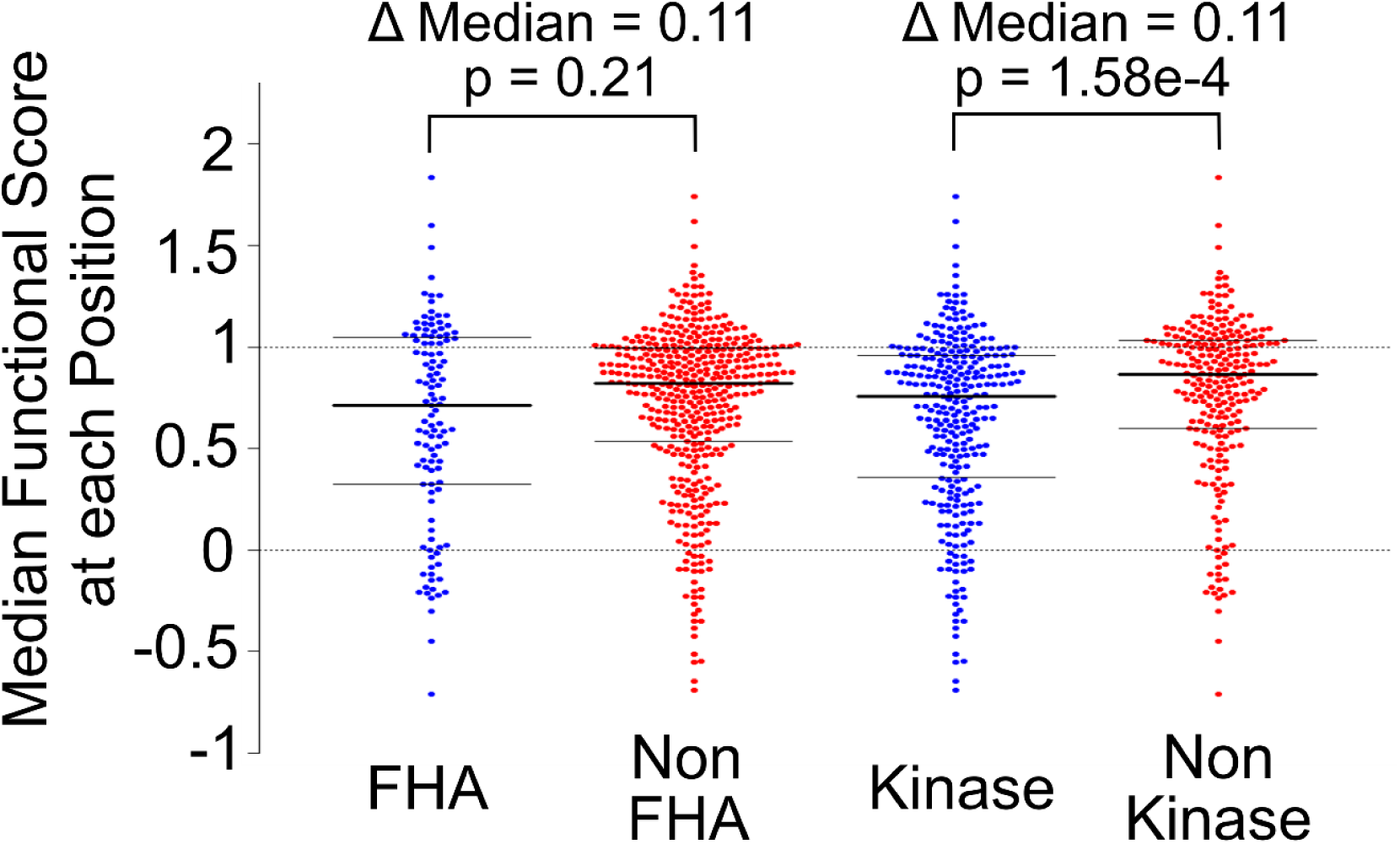
Comparing Functional scores in the FHA and kinase domain to the rest of CHK2. The median functional score for missense variants at each position were stratified into those located in the FHA domain (92 to 205), the kinase domain (212 to 501), and those not located in the FHA domain (2 to 91 and 206 to 543) or not in the kinase domain (2 to 211 and 502 to 543). Positions located in the FHA domain were compared to those outside the FHA, as well as kinase domain positions compared to non-kinase domain, by Wilcoxon rank-sum test. The median value and 25th and 75th quantiles are overlaid on each distribution as solid horizontal lines. The dashed horizontal lines across all distributions indicate scores of zero and one, indicating deleterious and neutral variants respectively.

**Figure S8.**
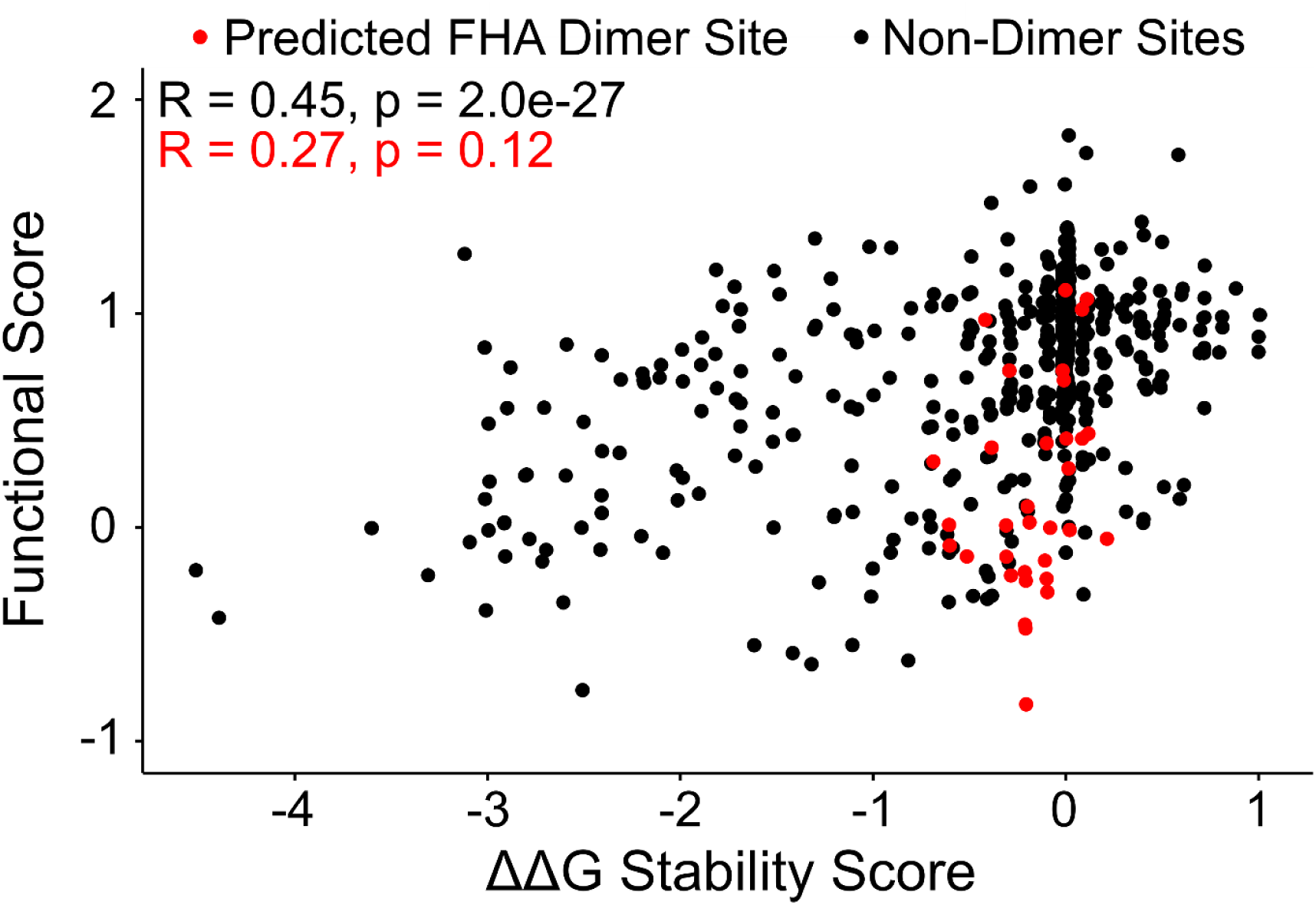
Position-specific ΔΔG stability and functional scores for CHK2. The median ΔΔG score and median functional score was calculated for each position along CHK2 and depicted as a scatterplot. Positions with ΔΔG scores near and below - 1 are predicted to impact protein stability while scores of 0 are not predicted to destabilise the protein. Functional scores near 1 are functionally neutral while scores 0 and below are deleterious. The proposed FHA dimerization sites (93 to 97, 115 to 117, 138 to 144, 164 to 169, and 191 to 202) are highlighted in red with the rest of the protein shown in black. Pearson correlations for proposed dimer sites are also shown in red alongside Pearson correlation for the rest of *CHK2*.

**Figure S9.**
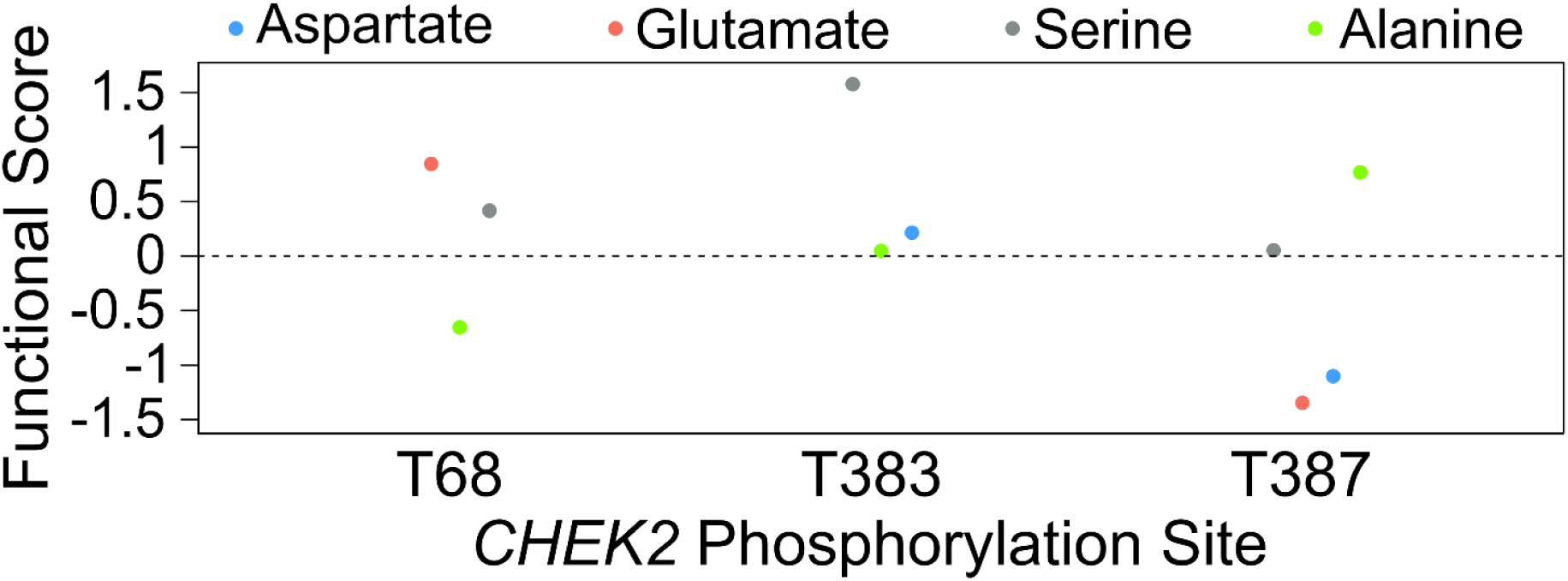
Effect of phosphomimetic and phosphodead variants at known CHK2 phosphorylation sites. Functional scores are shown for individual variants located at phosphorylation sites critical for CHK2 activation. Phosphomimetic mutations, aspartate and glutamate, are shown in blue and red respectively, the purportedly neutral variant (due to its ability to be phosphorylated) serine is shown in grey, and phosphodead alanine shown in green.

**Figure S10.**
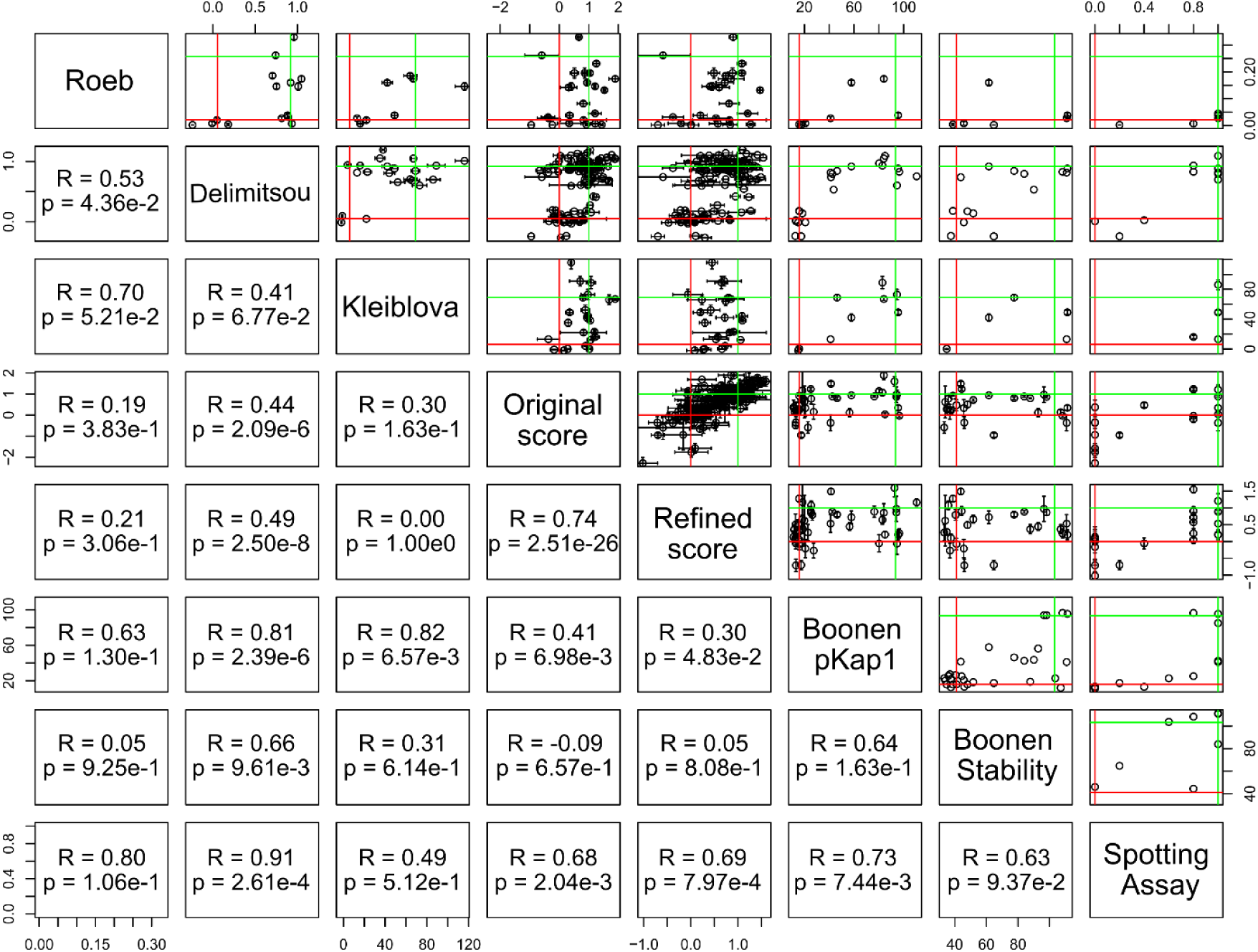
Scatterplots comparing all functional assays. Scatterplots were generated comparing each of the functional assay datasets indicated on the diagonal axis. Starting from the top-left, this includes Roeb *et al.* 2012, Delimitsou *et al.* 2019, Kleiblova *et al.* 2019, our experimental and refined functional scores, Boonen *et al.* 2022 pKAP1 and stability scores, and finally MMS spotting assays we performed for individual variants (data not shown). Correlation between datasets was assessed by Pearson coefficient.

**Figure S11.**
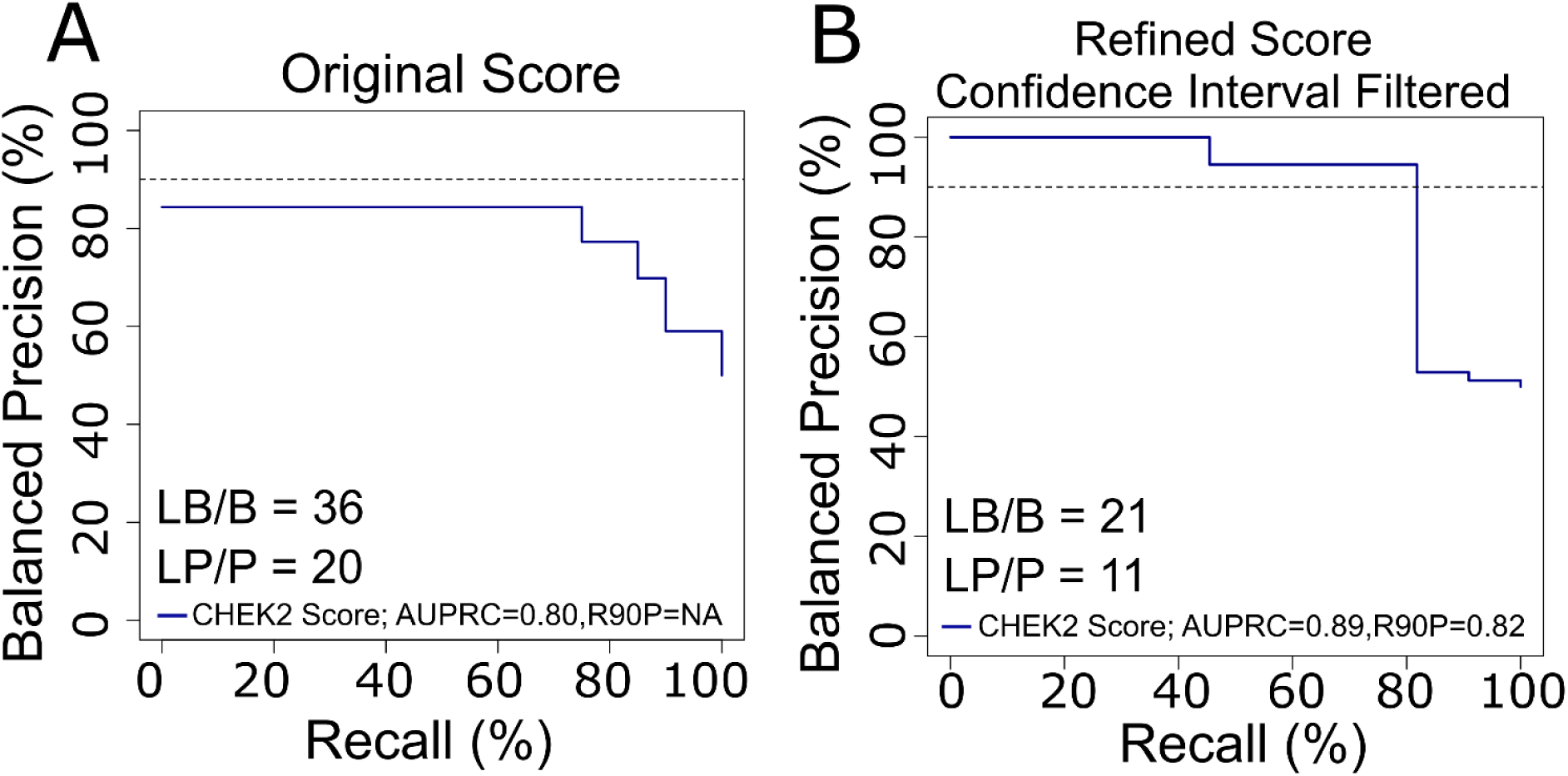
Balanced Precision-Recall Curves of functional scores against variant with known pathogenic or benign annotations. **A** Using the direct original *CHEK2* map scores and a known set of clinically annotated pathogenic or benign *CHEK2* variants from Invitae, we evaluated balanced precision—defined at each score threshold by the fraction of variants that are pathogenic given a balanced (50% prior probability of pathogenicity) test set—versus recall (fraction of pathogenic variants captured at this threshold). The horizontal dashed line indicates R90BP with the numerical AUPRC and R90BP listed in the bottom-left hand legend. LB/B indicates likely benign and benign while LP/P means likely pathogenic and pathogenic. **B** A balanced precision-recall curve was generated as described above for the imputed and refined functional scores after confidence interval filtering.

